# Defining the KRAS-regulated kinome in KRAS-mutant pancreatic cancer

**DOI:** 10.1101/2021.04.27.441678

**Authors:** J. Nathaniel Diehl, Jennifer E. Klomp, Kayla R. Snare, Devon R. Blake, Priya S. Hibshman, Zane D. Kaiser, Thomas S.K. Gilbert, Elisa Baldelli, Mariaelena Pierobon, Björn Papke, Runying Yang, Richard G. Hodge, Naim U. Rashid, Emanuel F. Petricoin, Laura E. Herring, Lee M. Graves, Adrienne D. Cox, Channing J. Der

## Abstract

Oncogenic KRAS drives cancer growth by activating diverse signaling networks, not all of which have been fully delineated. We set out to establish a system-wide profile of the KRAS-regulated kinase signaling network (kinome) in KRAS-mutant pancreatic ductal adenocarcinoma (PDAC). We knocked down KRAS expression in a panel of six cell lines, and then applied Multiplexed Inhibitor Bead/Mass Spectrometry (MIB/MS) chemical proteomics to monitor changes in kinase activity and/or expression. We hypothesized that depletion of KRAS would result in downregulation of kinases required for KRAS-mediated transforming activities, and in upregulation of other kinases that could potentially compensate for the deleterious consequences of the loss of KRAS. We identified 15 upregulated and 13 downregulated kinases in common across the panel. In agreement with our hypothesis, all 15 of the upregulated kinases have established roles as cancer drivers (e.g., SRC, TGFBR1, ILK), and pharmacologic inhibition of the upregulated kinase, DDR1, suppressed PDAC growth. Interestingly, 11 of the 13 downregulated kinases have established driver roles in cell cycle progression, particularly in mitosis (e.g., WEE1, Aurora A, PLK1). Consistent with a crucial role for the downregulated kinases in promoting KRAS-driven proliferation, we found that pharmacologic inhibition of WEE1 also suppressed PDAC growth. The unexpected paradoxical activation of ERK upon WEE1 inhibition led us to inhibit both WEE1 and ERK concurrently, which caused further potent growth suppression and enhanced apoptotic death than WEE1 inhibition alone. We conclude that system-wide delineation of the KRAS-regulated kinome can identify potential therapeutic targets for KRAS-mutant pancreatic cancer.

## Introduction

Pancreatic ductal adenocarcinoma (PDAC) is among the deadliest cancers, with a 5-year survival rate of 10% (1). Despite significant advances in our understanding of the genetic and molecular drivers of PDAC (2), effective targeted therapies are still lacking. Current standards-of-care are comprised of conventional cytotoxic drugs (*3*).

Mutational activation of the *KRAS* oncogene is the initiating genetic event and is found in 95% of PDAC (4). There is now substantial experimental evidence that KRAS is essential for the maintenance of PDAC growth, and consequently, the National Cancer Institute has identified the development of anti-KRAS therapeutic strategies as one of four major initiatives for the field (5). Recent early-stage clinical findings with direct inhibitors of one KRAS mutant, G12C, support the potential clinical impact of an effective KRAS inhibitor (6, 7). However, while KRAS^G12C^ mutations are common in lung adenocarcinoma, they comprise only 2% of *KRAS* mutations in PDAC (4, 8). Therefore, indirect strategies to block aberrant KRAS signaling remain arguably the best approach for the majority of KRAS-mutant PDAC (9).

One key approach is to inhibit downstream effector signaling. Of the multitude of KRAS effectors, at least four are validated drivers of KRAS-dependent PDAC growth (8). The best characterized and potentially most crucial is the RAF-MEK-ERK mitogen-activated protein kinase (MAPK) cascade. That activated *Braf* can phenocopy mutant *Kras* and, together with *Tp53* mutations, drive full development of metastatic PDAC in mouse models supports the key role of the MAPK cascade in driving Kras-dependent PDAC growth (10). While the substrates of RAF and MEK kinases are highly restricted, ERK1/2 serine/threonine kinase activation can cause direct or indirect phosphorylation of a diverse spectrum of more than one thousand proteins (11). Since ERK substrates include other protein kinases (e.g., RSK1-4, MNK2 and MSK1/2), ERK activation can regulate a highly diverse phosphoproteome (12). However, the specific components are context-dependent, and the ERK-regulated phosphoproteome downstream of KRAS in PDAC is not well delineated.

The second best characterized KRAS effector are phosphoinositide 3-kinases (PI3K) (13). KRAS activation of PI3K promotes formation of phosphatidylinositol (3,4,5)-triphosphate, leading to activation of the AKT and mTOR serine/threonine kinases. Activation of PI3K-AKT-mTOR signaling has also been observed as a compensatory response to ERK inhibition, driving resistance to ERK MAPK inhibitors (14, 15). Thus, concurrent PI3K inhibition synergistically enhances the anti-tumor activity of ERK MAPK inhibitors (10). Other key KRAS effectors include the RAL and RAC small GTPases, that activate TBK1 and PAK serine/threonine kinases (8). With their high tractability as therapeutic targets, these and other protein kinase components of KRAS effector signaling have been pursued as indirect approaches to KRAS inhibition (16). We propose that still other kinases may also be of interest as targets in KRAS-mutant PDAC.

Additionally, a limitation of essentially all targeted anti-cancer therapies is the onset of treatment-induced acquired resistance (17, 18). In response to pharmacologic inhibition of a key cancer driver, cancer cells can induce a complex array of responses that functionally compensate for target inhibition. One major mechanism of resistance is loss of the negative feedback signaling that tempers the strength of growth regulatory signaling networks (19). For example, although aberrant ERK activation can drive cancer growth, excessive ERK activity is deleterious, causing growth cessation through induction of senescence or apoptosis (20–22). Therefore, a multitude of ERK-dependent negative feedback loops exist to dampen ERK activity (23). Thus, upon pharmacological inhibition of ERK, loss of ERK-dependent negative feedback leads to ERK reactivation to overcome inhibitor action. Similar responses are also seen upon inhibition of the PI3K-AKT-mTOR pathway (24).

An important strategy to improve the long-term efficacy of targeted therapies is the elucidation of treatment-induced resistance mechanisms, which can then be used to identify drug combinations capable of blocking or even reversing the onset of resistance. Unbiased system-wide screening methods applied for this purpose include powerful genetic approaches such as CRISPR- or RNAi-mediated loss-of-function or cDNA overexpression/activation gain-of-function screens (25). For example, a CRISPR-Cas9 screen demonstrated that ERK reactivation is a primary mechanism limiting the efficacy of KRAS^G12C^ inhibitors (26). These findings have guided initiation of ongoing clinical trials to evaluate KRAS^G12C^ inhibitors in combination with inhibitors that act either upstream (e.g., on EGFR) or downstream (e.g., on MEK) of KRAS (clinicaltrials.gov). Similarly, Wood et al. applied an activated signaling expression library to identify both known and novel mechanisms that drive resistance to ERK MAPK inhibitors (27). In a complementary approach, our chemical library screen demonstrated that concurrent inhibition of other ERK MAPK components, to block compensatory ERK reactivation, synergistically enhanced the efficacy of RAF inhibitors in KRAS-mutant PDAC (28).

In light of the key role of kinome reprogramming in driving resistance, it is crucial to be able to detect changes in kinase activity and/or expression in response to pharmacologic inhibition of an oncogenic signaling driver. Chemical proteomics is a particularly powerful experimental technique that enables kinome-wide profiling of these changes. One version of this type of assay, MIB/MS, incorporates multiplexed inhibitor beads (MIBs), coupled to mass spectrometry (MS) (29). MIBs, comprised of broad-spectrum kinase inhibitors covalently coupled to Sepharose beads, can be used to monitor protein kinase expression and/or activation, where MIB-associated kinases are identified by MS. In our initial application of MIB/MS, we identified activation of multiple receptor tyrosine kinases (RTKs) in response to MEK inhibitor (MEKi) treatment that may drive MEKi resistance in KRAS-mutant breast cancer (30). Accordingly, concurrent inhibition of RTKs and MEK more effectively impaired transformed and tumor growth in vitro and in vivo than MEKi alone. We have also applied MIB/MS to identify novel inhibitors of chemotherapy-resistant PDAC (31), to pinpoint tumor cell-type specific responses to the clinical kinase inhibitor dasatinib in diffuse large B cell lymphoma (32), and to identify EGFR/HER2 activation of the MEK5-ERK5 MAPK cascade as compensatory response to ERK1/2 inhibition, which drives resistance to ERK inhibitor treatment by stabilizing the MYC oncoprotein (33).

In the present study, we hypothesized that a system-wide delineation of KRAS-regulated kinases can identify therapeutic targets that may not have been considered based on our current knowledge of KRAS effector signaling networks. We speculated that KRAS may require additional kinases beyond the classical RAF-MEK-ERK and PI3K-AKT-mTOR effector pathways to promote its transforming functions, and that these would be downregulated upon KRAS depletion. In contrast, we anticipated that yet other kinases would be upregulated as compensatory responses to this depletion. Thus, identification of both upregulated as well as downregulated kinases may establish novel targets for anti-KRAS therapies. We therefore applied the kinome-wide MIB/MS assay to elucidate an unbiased profile of the KRAS-dependent kinome in PDAC. Our strategy further elucidates the complex spectrum of protein kinases functionally linked to aberrant KRAS activation and identifies unanticipated signaling vulnerabilities and potential therapeutic approaches for PDAC.

## Results

### KRAS suppression alters the activity of diverse kinases in PDAC

Although oncogenic KRAS effector signaling activates the ERK MAPK cascade and other protein kinases, we speculated that the full spectrum of KRAS-regulated protein kinases remained to be elucidated. To gain a full understanding of the KRAS-dependent kinome in KRAS-mutant PDAC, we suppressed *KRAS* expression in a panel of six KRAS-mutant PDAC cell lines using validated short interfering RNA (siRNA) (Fig. S1*A* and data file S1) (14). After 72 h, a time point at which compensatory changes in the kinome have been initiated in response to loss of KRAS (28), cell lysates were processed for MIB/MS label-free proteomics to monitor kinome-wide changes in activity and/or expression (Fig. S1*B*). Our approach detected a total of 227 kinases of sufficient abundance for quantification in one or more PDAC cell lines (Fig. S1*C*). As expected, given the genetic heterogeneity of PDAC tumors, there was significant heterogeneity in the kinase profile of each cell line (Fig. S1*C*). Interestingly, the quantified kinases were upregulated and downregulated to the same degree (63 and 62 kinases, respectively; Fig. S1*D*). Across all six lines, we identified 15 kinases that were significantly upregulated and 13 that were downregulated in common (Fig. 1*A* and Fig. S1*E*).

**Figure. 1.**
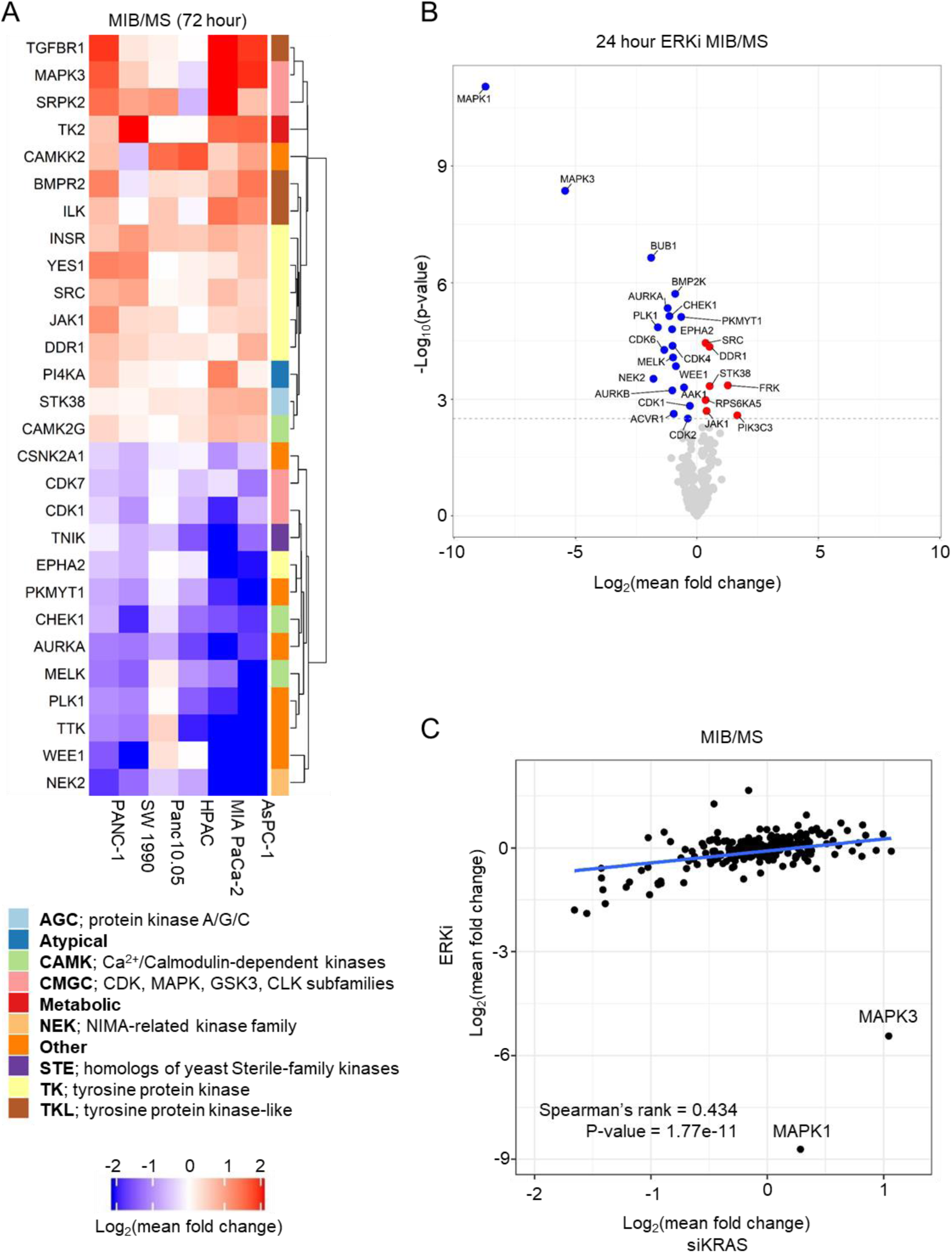
KRAS knockdown induces kinome-wide alterations primarily through ERK. *A*, heatmap summarizing kinases that were significantly altered (p adjusted < 0.05) after 72 h of non-specific control siRNA (siNS) or siKRAS treatment, followed by MIB/MS enrichment. Lysates were collected from six KRAS-mutant PDAC cell lines [SW 1990, MIA PaCa-2, PANC-1, HPAC, AsPC-1 and Panc 10.05]. Unsupervised clustering was used to visualize distance metrics of log-transformed fold change values for significant kinases following ANOVA analysis. Kinase class annotations were added from UniProt classification and index of kinases. *B*, volcano plot showing results of significance testing after ANOVA [y-axis] with the log-transformed mean fold change in ERKi (SCH772984) over DMSO control (24 h). Significant kinases are colored red [increased] or blue [decreased] if p-value is significant (p.adj < 0.05) following Benjamini-Hochberg correction (n = 3). *C*, Spearman’s rank correlation analysis of siKRAS/siNS samples versus ERKi/DMSO samples, 222 kinases were shared among dataset.

To determine if the deregulated kinases represented specific subgroups, we applied the UniProt kinase classification annotation that is based in part on catalytic domain sequence identity and in part on biological function (Fig. S1*F*). Notably, although the downregulated kinases did not share much sequence identity (7/13 or 53.8% were “Other” kinases versus 32/227 or 14.1% of all detected kinases) (Fig. S1, *F* and *G*), nearly all have established roles in regulating cell cycle transitions, particularly through mitosis (e.g., Aurora kinase A (*AURKA*), CHK1 (*CHEK1*), WEE1) (table S1).

In contrast, the upregulated kinases were enriched in tyrosine kinases (TK) and tyrosine-like kinases (TKL) (7/15; 46.7%), which comprised 23.3% (53/227) of all detected kinases (Fig. S1, *1* and *H*). Strikingly, a driver role in cancer has been attributed to all 15 upregulated kinases (e.g., ERK1 (*MAPK3*) (table S2), SRC and JAK), supporting the strong likelihood that upregulated kinases serve compensatory growth-promoting roles that attenuate the deleterious consequences of KRAS deficiency. Supporting this possibility, we showed recently that ERK inhibitor treatment of PDAC cell lines caused upregulation of SRC activity and that concurrent SRC inhibition further enhanced ERK inhibitor-mediated growth suppression (33).

To complement these kinome analyses, we used reverse phase protein array (RPPA) pathway activation mapping (34) to evaluate the signaling consequences of 72 h of siRNA-mediated *KRAS* suppression (Fig. S1*I*). The RPPA panel included 149 phospho-specific and total antibodies to monitor the activation state or expression, respectively, of signaling proteins involved in regulation of cell proliferation, survival, motility, etc. (table S3 and data file S2). As expected, phosphorylated and total ERK proteins were increased, consistent with loss of ERK-dependent negative feedback signaling (Fig. S1*J*). Nevertheless, phosphorylation of key ERK substrates important for driving ERK-dependent growth (RSK, ELK1 and MYC) remained strongly suppressed (Fig. S1*J*). Thus, similar to our previous observations with pharmacologic inhibition of ERK (14, 15), the level of ERK phosphorylation did not reliably reflect the level of ERK signaling.

Among the decreases in expression and/or activation of multiple proteins involved in cell cycle regulation were a dramatic reduction in the mitotic marker, phosphorylated histone H3 at Ser^10^, as well as loss of the proliferation markers Ki67 and phosphorylation of RB at Ser^780^ (Fig. S1*I*). RPPA analyses also revealed alterations in expression (e.g., BIM) or phosphorylation (e.g., FADD Ser^194^) of proteins associated with apoptosis, a well-described consequence of loss of mutant KRAS function (35). Finally, reductions were also detected in phosphorylated S6 ribosomal protein, involved in regulation of protein translation (Fig. S1, *1* and *J*). Thus, KRAS loss is associated with the activation and inactivation of diverse growth regulatory signaling pathways.

We next wanted to determine the contribution of the ERK MAPK effector pathway to the regulation of the KRAS-dependent kinome. We treated the same panel of six PDAC cell lines for 24 h with the ERK1/2-selective inhibitor SCH772984 (ERKi) (36), then performed MIB/MS analyses (data file S3). Quantification of kinases revealed that, of 264 kinases detected, 26 were significantly altered in expression and/or activity across all six cell lines (Fig. 1*B* and Fig. S1*K*). Of these, seven were upregulated and 19 were downregulated. As expected, ERK1 and ERK2 (*MAPK3* and *MAPK1*) activities were by far the most strongly downregulated upon ERKi treatment (Fig. 1*B*).

Importantly, many of the same kinases were altered upon pharmacological inhibition of ERK and upon genetic suppression of KRAS. Nine of the kinases downregulated by siKRAS (Fig. 1*A*) were also downregulated by ERKi (Fig. 1*B*), including many involved in progression through mitosis. Similarly, DDR1, JAK1 and SRC were among kinases upregulated by both siKRAS and ERKi. Spearman’s rank correlation analysis of siKRAS- and ERKi-mediated kinome changes revealed significant overlap between the 222 kinases detected in both datasets (rho = 0.434, p = 1.77e-11) (Fig. 1*C*). The significant overlap in kinase signaling changes following either ERK inhibition or KRAS suppression is consistent with the predominant role of ERK MAPK in supporting KRAS-dependent PDAC growth (10, 14). We conclude that the observed kinome changes conferred by genetic suppression of KRAS were mediated in large part through loss of ERK1/2 signaling.

### Inhibition of DDR1, but not JAK, impairs PDAC growth

We speculated that upregulated kinases, in particular those with known driver roles in cancer, may represent compensatory activities in response to loss of KRAS. Among the kinases upregulated following KRAS knockdown, JAK1 was notable because the JAK1-STAT3 signaling axis has been shown to be important for PDAC tumorigenesis and can drive resistance to inhibition of MEK (37, 38). Consistent with MIB/MS, RPPA also showed that four of five cell lines displayed significant (p-adjusted < 0.05) activation of STAT3 following KRAS knockdown (Fig. 2*A*), as indicated by phosphorylation at the JAK family site Tyr^705^ (39). Phosphorylation of STAT3 at Ser^727^, which is not a JAK site but a possible ERK or JNK target (40), also increased in a cell line dependent manner (40), as did phosphorylation of STAT1 and STAT5. STAT6 phosphorylation at Tyr^641^ was the most consistent of all, in agreement with our previous finding of significant upregulation of JAK signaling through this site upon blockade of the MAPK cascade (28). To determine whether JAK1-STAT3 was similarly affected, we directly inhibited signaling through ERK MAPK. In the presence of the ERK1/2 inhibitor SCH772984, STAT3 phosphorylation (Tyr^705^) was increased in Pa02C and Pa16C and unchanged in MIA PaCa-2 and PANC-1 cells (Fig. 2*B*). We conclude that upregulation of JAK1-STAT signaling induced by KRAS suppression was mediated through loss of ERK effector signaling.

**Figure 2.**
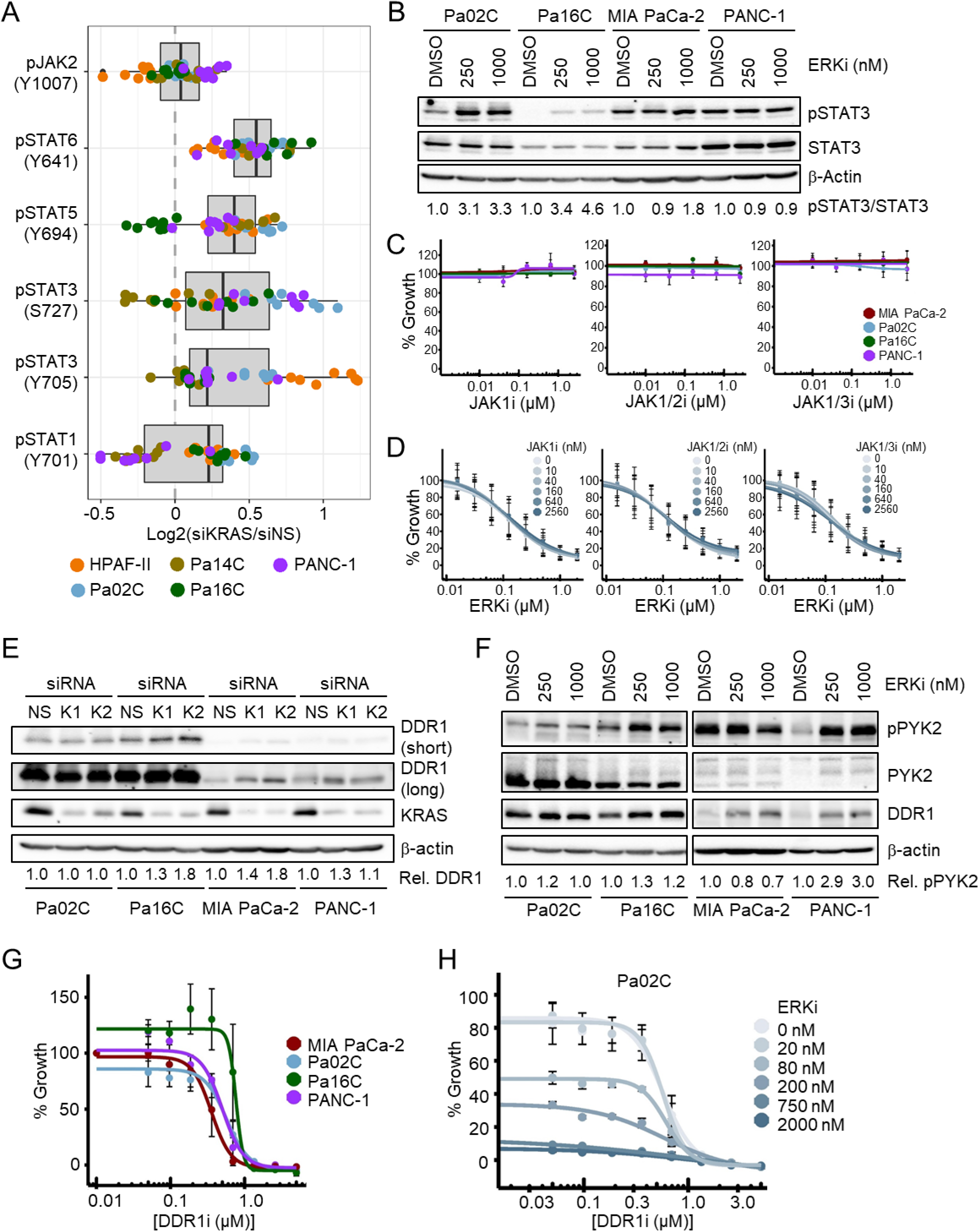
DDR1, but not JAK, is a therapeutic vulnerability in PDAC. *A*, reverse phase protein array (RPPA) of five PDAC cell lines following 72 h siRNA knockdown of KRAS. Values are log_2_-transformed fold change with respect to median siNS value (n = 4). *B*, immunoblot of PDAC cell lines following treatment with vehicle or ERKi (SCH772984) for 24 h (n = 4). Ratios of pSTAT3/STAT3 are reported below the blot. *C*, proliferation of PDAC cells (5 days) following treatment with JAK1i (filgotinib), JAK1/2i (ruxolitinib), or JAK1/3i (tofacitinib) at various concentrations. Cell numbers at endpoint were normalized to vehicle-treated control (100% growth) for each cell line. Curves were fit using a 4-parameter log-logistic function and the drm package in R (n = 3). *D*, Proliferation of Pa02C cells (5 days) following combination treatment with JAK1i, JAK1/2i, or JAK1/3i and ERKi (SCH772984) at various concentrations. Cell numbers were normalized and curves were fit as in panel *C* (n = 3). *E*, immunoblot of PDAC cell lines following treatment with non-specific (NS) or KRAS-targeting (K1, K2) siRNA constructs for 72 h. Relative DDR1 levels are indicated below the blot (total DDR1 normalized for loading differences as determined by β-actin). “Short” and “long” indicate shorter and longer exposures of the same membrane, respectively (n = 3). *F*, Immunoblot of PDAC cell lines following treatment with vehicle or ERKi (SCH772984) for 24 h (n = 4). Relative pPYK2 levels (vs. DMSO) in each cell line are indicated below the blot. *G*, Proliferation of PDAC cells (5 days) following treatment with DDR1i (7rh) at various concentrations. Cell numbers were normalized and curves were fit as in panel *C* (n = 3). *H*, Proliferation of Pa02C cells (5 days) following combination treatment with DDR1i (7rh) and ERKi (SCH772984) at various concentrations. Cell numbers were normalized and curves were fit as in panel (*C*) (n = 3).

We next determined if upregulation of JAK signaling is a compensatory response that can offset the growth suppression induced by loss of KRAS or ERK. We utilized three different JAK inhibitors, targeting JAK1 (filgotinib), JAK1/2 (ruxolitinib), and JAK1/3 (tofacitinib) (41). All three JAK inhibitors reduced STAT3 phosphorylation in a dose-dependent manner, though PANC-1 cells were resistant to filgotinib (Fig. S2*A*). JAK inhibitors alone had no effect on anchorage-dependent proliferation of four PDAC cell lines (Fig. 2*C*), indicating the lack of dependence on JAK when KRAS is present. Further, co-treatment of four PDAC cell lines with each JAK inhibitor in combination with ERKi revealed that concurrent JAK inhibition did not further enhance ERKi-mediated growth suppression (Fig. 2*D* and Fig. S2*B*). Thus, while JAK activity was increased upon loss of KRAS, subsequent inhibition of JAK family kinases did not potentiate the antiproliferative effect of direct ERK inhibition in PDAC.

To further investigate whether upregulated kinases could compensate for the loss of KRAS, we next evaluated the discoidin domain receptor 1 (DDR1). DDR1 is a receptor tyrosine kinase that transduces collagen-mediated proliferative signaling from stroma-associated extracellular matrix (42), and pharmacologic inhibition of DDR1 function impaired PDAC growth in vivo in a mouse model (43, 44). Given our identification of DDR1 as a kinase that was upregulated upon KRAS suppression in PDAC cell lines in vitro (Fig. 1*A*), we sought to determine whether upregulated DDR1 could also serve as an increased pro-proliferative signal in a KRAS-dependent, tumor cell-intrinsic manner. We first determined that both KRAS knockdown and ERK inhibition increased DDR1 protein levels (Fig. 2, *E* and *F*, respectively). Additionally, ERK inhibition increased phosphorylation of the DDR1 effector PYK2 (pPYK2, Fig. 2*F*) at the DDR1 phosphorylation site, Tyr^402^ (44), in three of four cell lines. Thus, we confirmed that DDR1 signaling is responsive to KRAS-ERK signaling.

To evaluate the possibility that DDR1 upregulation is a compensatory response to the growth suppression mediated by loss of KRAS-ERK, we first determined if inhibition of DDR1 alone is deleterious to growth. We treated PDAC cells with 7rh, a potent and selective ATP-competitive DDR1 inhibitor (45) and observed comparable IC_50_ (suppression of PYK2 Tyr^402^) and GI_50_ values, supporting on-target growth inhibition (Fig. 2*G* and Fig. S2, *2* and *D*) (45). We found that concurrent inhibition of DDR1 and ERK caused dose-dependent loss of cell viability (Fig. 2*H* and Fig. S2*E*), ranging between additivity and synergy, according to BLISS analysis (Fig. S2*F*). Overall, our evaluation of kinases upregulated by siKRAS or ERKi has revealed a novel compensatory mechanism in DDR1 and a surprising dispensability of JAK signaling for PDAC anchorage-dependent cell growth and viability.

### Loss of KRAS or MAPK signaling causes loss of G2/M and DNA-damage response kinases

In order to better understand the relationships among downregulated kinases following KRAS genetic suppression, we performed STRING analysis on significantly altered kinases (Fig. 3*A*). Eleven of the 13 downregulated kinases formed a tight interaction node, with only EPHA2 and TNIK involved in distinctly separate signaling networks. Next, we utilized Panther Gene Ontology to determine the relevant biological processes (Fig. 3*B*). Given that our queried genes exclusively encode protein kinases, it was expected that “protein phosphorylation” was the top gene set identified. Additionally, a main function of KRAS signaling through the ERK cascade is to promote cell cycle progression through G1 by transcriptional stimulation of *CCND1* (encoding cyclin D1), leading to CDK4/6-dependent phosphorylation and inactivation of the RB tumor suppressor (46). Thus, it was unexpected that we observed that multiple downregulated kinases are involved in G2/M checkpoint and mitotic functions (Table S1). Consistent with KRAS promotion of G1/S cell cycle progression, flow cytometry analyses in four PDAC cell lines showed that KRAS knockdown caused an increase (14-24%) in cells in G1 and a corresponding decrease (15-22%) in cells in S, albeit no significant change in the G2/M cell population (Fig. S3, *3* and *B*).

**Figure 3.**
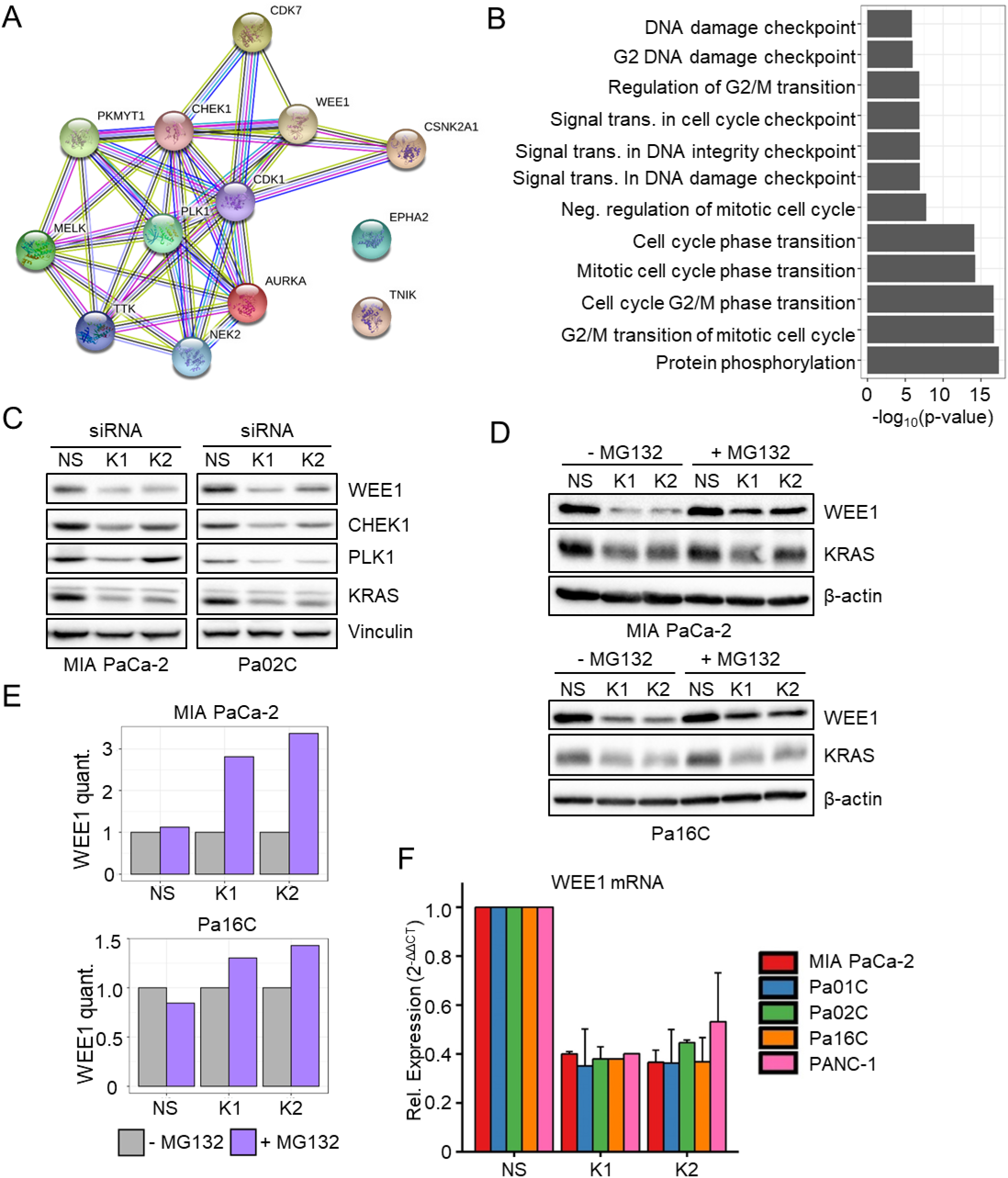
Loss of KRAS/MAPK signaling contributes to loss of G2/M and DNA-damage response kinases. *A*, downregulated kinase identifiers from the MIB-MS screen (Fig. 1A) were used as input into the STRING database, and the resulting bitmap was downloaded. Edges (lines between nodes) represent known or predicted protein-protein associations. *B*, downregulated kinase identifiers were used as input for Gene Ontology enrichment analysis of biological processes (72, 73). Selected enriched gene sets (p-adjusted < 0.05) are displayed along the y-axis, with the –log_10_-transformed p-values displayed along the x-axis. (**C**) immunoblot analysis of PDAC cell lines following treatment with non-specific (NS) or KRAS-targeting (K1, K2) siRNA constructs for 72 h (n = 4). *D*, Protein stability of WEE1 as determined by 4 h MG132 proteasomal inhibitor treatment following 26 h treatment with non-specific (NS) or KRAS-targeted (K1, K2) siRNA constructs (n = 3). *E*, Quantification of blot in panel *D*. Densitometry estimates were first normalized for loading efficiency using β-actin. Next, each sample value was calculated as a ratio of MG132+/MG132- and paired by siRNA construct. *F*, qRT-PCR was performed on five PDAC cell lines following 72 h treatment with non-specific (NS) or KRAS-targeted (K1, K2) siRNA. Relative expression [y-axis] of WEE1 transcripts was measured (n = 3).

Interestingly, the 13 downregulated kinases identified in the chemical proteomics screen were also enriched for roles in DNA damage checkpoints and maintaining DNA integrity (Fig. 3*B*). We therefore queried our RPPA data for genes related to both DNA-damage repair and cell cycle, which revealed a mixed response to KRAS knockdown (Fig. S3*C*). Cyclin D1 levels were reduced whereas cyclin A levels either increased or remained unchanged. Additionally, phosphorylation of ATM at Ser^1981^ increased whereas phosphorylation of ATR at Ser^428^ was diminished in all cell lines. Taken together, these changes suggest that loss of KRAS causes a loss of specific cell cycle promoting factors. Additionally, the changes in ATM and ATR phosphorylation, though divergent, suggest that KRAS knockdown may induce genomic stress that could also contribute to cell cycle factor changes.

While significant literature points to the importance of KRAS and ERK in promoting G1 progression, fewer studies outline the role of mutant KRAS in maintaining DNA damage checkpoint kinases such as WEE1, PKMYT1, and CHEK1 (Fig. 3*A*) (47–49). We confirmed that KRAS knockdown caused loss of WEE1, CHEK1, and PLK1 proteins (Fig. 3*C*). A recent transcriptome analysis of PDAC identified WEE1 as a potential therapeutic target (50). Therefore, we focused our further investigations on this DNA-damage checkpoint kinase.

To address a basis for the loss of WEE1 protein caused by KRAS suppression, we first examined WEE1 protein stability. Treatment with the proteasome inhibitor MG132 prevented WEE1 degradation more robustly in KRAS-suppressed samples as compared to control samples. This suggests that the absence of KRAS signaling destabilizes WEE1 protein (Fig. 3, *D* and *E*, and Fig. S3, *3* and *E*). We hypothesized that KRAS, through its extensive transcriptional network, may also regulate *WEE1* transcription. Indeed, RT-PCR analyses demonstrated that loss of KRAS decreased WEE1 transcripts in all five cell lines tested (Fig. 3*F*). We conclude that WEE1 protein stability and transcription are both reduced upon KRAS suppression.

### Inhibition of WEE1 kinase induces growth arrest and apoptosis in PDAC cells

The loss of WEE1 observed upon KRAS knockdown-induced growth suppression suggests that WEE1 activity may contribute to KRAS-mutant PDAC proliferation and therefore that direct inhibition of WEE1 may suppress KRAS-mediated cell proliferation. To address this possibility, we first treated cells with siRNA targeting *WEE1* and verified that loss of phosphorylation of the WEE1 substrate, CDK1 (Tyr^15^; pCDK1) was a reliable biomarker for loss of WEE1 function in PDAC (Fig. 4*A*). Similarly, treatment with the WEE1-selective clinical candidate inhibitor adavosertib/AZD1775 (WEE1i) dose-dependently reduced CDK1, with >80% reduction at 100 nM (Fig. 4*B* and Fig. S4*A*). Additionally, WEE1i treatment inhibited the proliferation of all six PDAC cell lines evaluated, with GI_50_ values (38 to 168 nM) comparable to the IC_50_ values (< 100 nM), supporting on-target growth suppression (Fig. 4*C* and Fig. S4, *A* to *C*). Similar growth inhibitory activities of WEE1i were also seen in clonogenic colony formation assays (Fig. 4*D*). Consistent with previous studies (51), we found that pharmacological inhibition of WEE1 led to accumulation in S- and G2/M-phases of the cell cycle (Fig. 4*E*, and Fig. S4, *4* and *E*). Additionally, WEE1i caused an approximately fivefold increase in apoptotic cells (Fig. 4*F*). Taken together, these results support our conclusion that downregulation of WEE1 contributes to the growth suppression induced by loss of KRAS.

**Figure 4.**
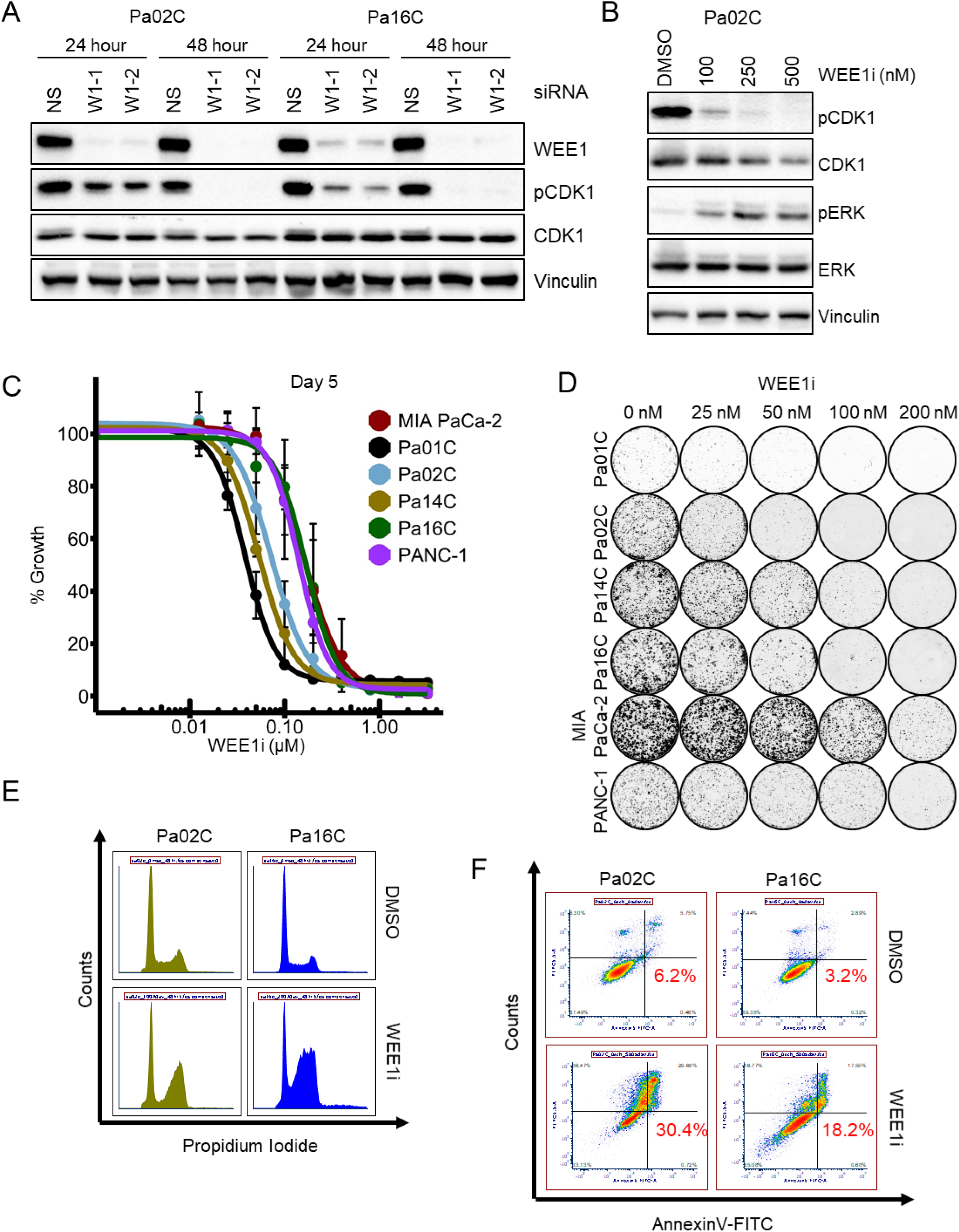
Inhibition of WEE1 kinase induces growth arrest and apoptosis in PDAC cells. *A*, Immunoblot analysis of PDAC cell lines following treatment with non-specific (NS) or WEE1-targeting (W1-1, W1-2) siRNA for 24 or 48 h (n = 3). *B*, immunoblot analysis of Pa02C cells following treatment with WEE1i for 48 h (n = 4). ***C***, proliferation of PDAC cells following treatment for 5 days with WEE1i (adavosertib) at various concentrations. Cell numbers at endpoint were normalized to vehicle-treated control (100% growth) for each cell line. Curves were fit using a 4-parameter log-logistic function and the drm package in R (n = 3). *D*, PDAC cells were plated at low density to allow for clonogenic growth and various concentrations of WEE1i were added. After 10 days, plates were stained with crystal violet and colonies were imaged (n = 4). *E*, cells were treated with vehicle or adavosertib (100 nM for Pa02C, 200 nM for Pa16C) for 24 h, then fixed and stained with propidium iodide. Flow cytometry quantification of cell cycle populations is shown for two representative cell lines (n = 3). *F*, annexin V-FITC staining followed by flow cytometry in PDAC cell lines after 72 h treatment with 500 nM WEE1i (n = 4).

### Combined inhibition of WEE1 and ERK causes synergistic growth arrest and apoptosis

Paradoxical activation of ERK has been reported previously in response to pharmacological inhibition of CHK1 (52, 53), another protein kinase involved in checkpoint inhibition in response to DNA damage. To determine if there is a similar response to WEE1 inhibition, we examined pERK in PDAC cell lines treated with adavosertib for 48 h. We observed a dose-dependent increase in pERK in all treated lines (Fig. 4*B* and Fig. S4, *4* and *F*). Evaluation at shorter time-points showed that WEE1 activity (pCDK1) was rapidly suppressed after 6 to 24 h, whereas increased pERK was seen only after 48 h (Fig. S4*C*). This delayed onset is consistent with compensatory reactivation of ERK to offset WEE1 inhibition-induced growth suppression. To address this possibility, we determined whether concurrent ERK inhibition enhances WEE1i growth inhibitory activity. We treated a panel of cell lines with both WEE1i and the ERK1/2- selective inhibitor SCH772984 (ERKi), which resulted in fewer colonies than treatment with either inhibitor alone (Fig. 5*A*). BLISS synergy analysis revealed that the combination exhibited synergistic activity in three cell lines (MIA PaCa-2, Pa16C and Pa02C), and additive activity in three cell lines (Fig. 5, *B* and *C*). Combining low-dose ERK inhibition (IC_25_-IC_50_) with WEE1 inhibition also synergistically induced apoptosis in four of four cell lines (Fig. 5, *D* and *E*). We conclude that concurrent inhibition of WEE1 and ERK synergistically induces both growth arrest and apoptosis in PDAC cell lines.

**Figure 5.**
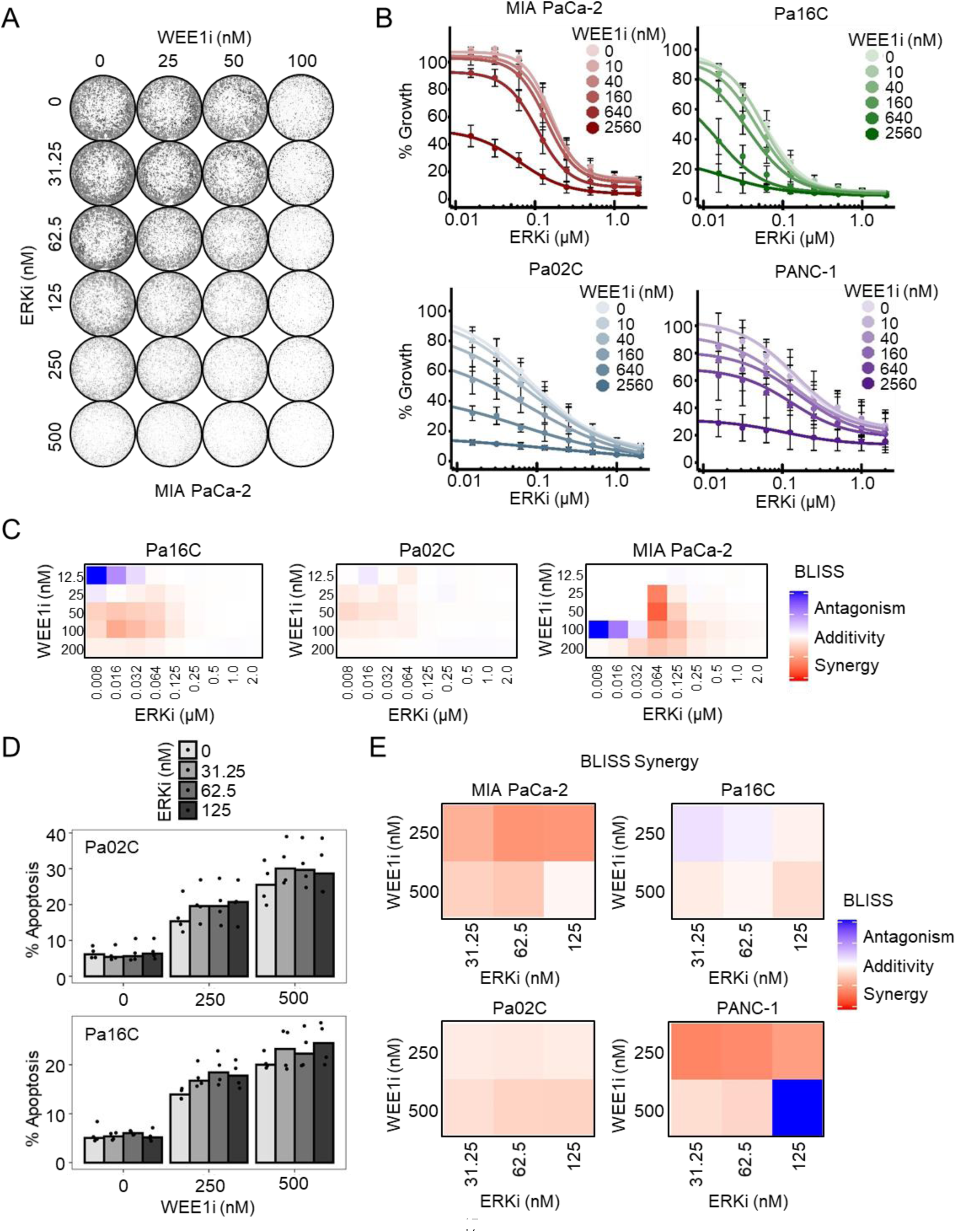
Combined inhibition of WEE1 and ERK induces growth arrest and apoptosis. *A*, PDAC cells were plated at low density and various concentrations of WEE1i (adavosertib) and/or ERKi (SCH772984) were added to the media. Cells were permitted to grow for 10 days to allow for clonogenic growth. Plates were then stained with crystal violet and imaged (n = 3). *B*, proliferation of PDAC cells following treatment for 5 days with WEE1i (adavosertib) and ERKi (SCH772984) at various concentrations in combination. Cell numbers at endpoint were normalized to vehicle-treated control (100% growth) for each cell line. Curves were fit using a 4-parameter log-logistic function and the drm package in R (n = 4). *C*, BLISS Synergy scores were calculated using the proliferation effect sizes from panel *B*. Scores < 1 indicate synergy (red), scores = 1 indicate additivity (white), and scores > 1 indicate antagonism (blue) (n = 4). *D*, annexin V-FITC staining followed by flow cytometry in PDAC cell lines after 72 h treatment with WEE1i. Quantification of apoptosis at each dose combination is displayed for the indicated cell lines (n = 4). *E*, BLISS Synergy scores were calculated using the proliferation effect sizes from panel *D*. Scores as for panel *C*, representative heatmaps are shown (n = 4).

### Inhibition of WEE1+ERK1/2 arrests growth of PDAC organoids

Three-dimensional (3D) patient-derived pancreatic cancer organoids maintained in Matrigel are believed to better model the therapeutic response of PDAC patients than cells grown in two-dimensional (2D) culture (54, 55). Therefore, we next evaluated the impact of concurrent WEE1 and ERK inhibition on proliferation of KRAS-mutant PDAC organoid cultures. The combination of inhibitors caused organoids to collapse and shrink more than either inhibitor alone (Fig. 6*A*, and Fig. S5, *5* and *B*). Concurrent WEE1 and ERK inhibition caused additive or synergistic effects on viability, depending on doses (Fig. 6, *B* and *C*). Taken together with anchorage-dependent growth assays, we conclude that concurrent ERKi treatment to block compensatory ERK activation can enhance WEE1 inhibition-mediated growth suppression.

**Figure 6.**
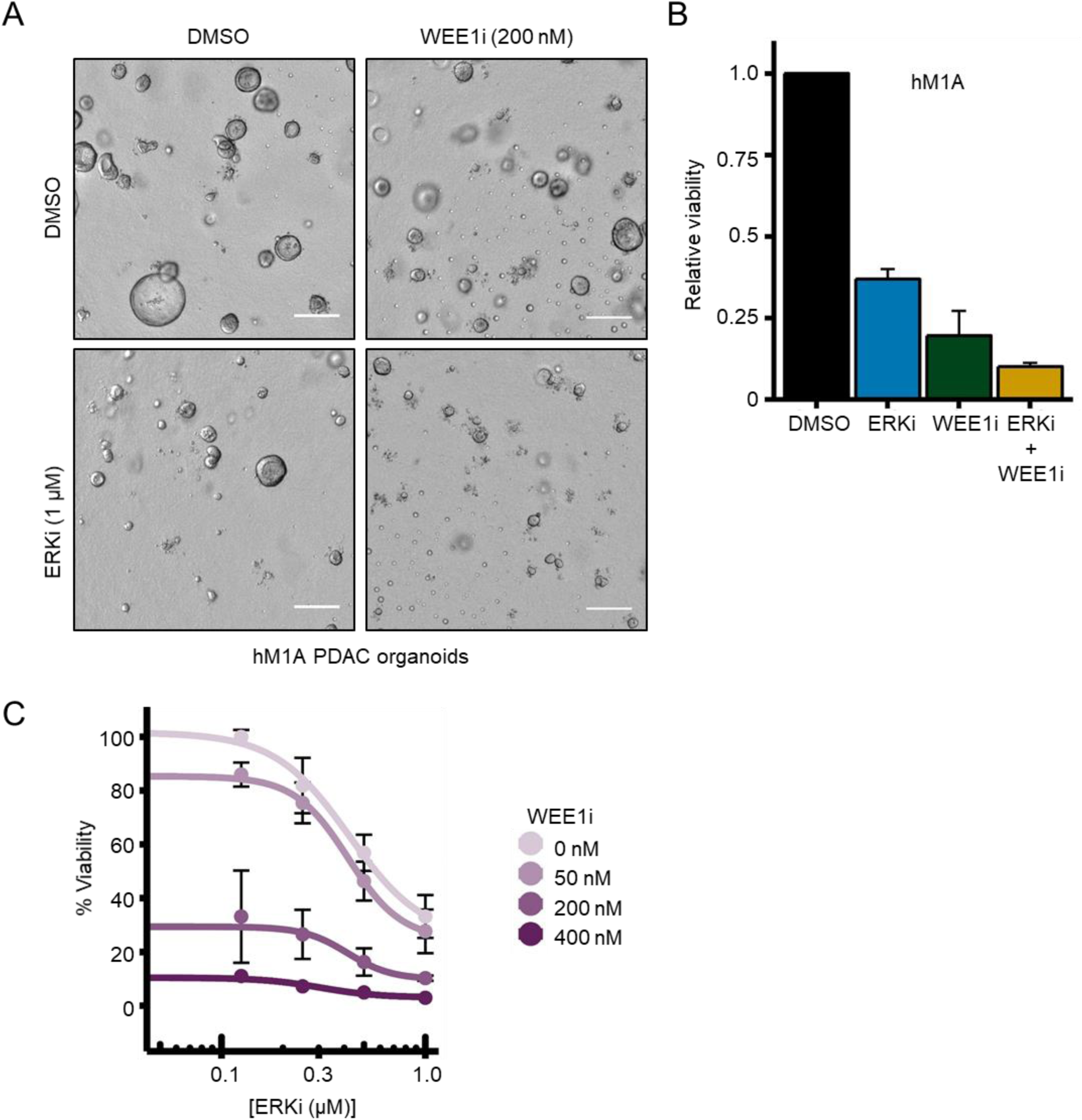
Inhibition of WEE1+ERK1/2 arrests growth of PDAC organoids. *A*, hM1A PDAC organoids were seeded for three days in Matrigel and organoid maintenance factors, followed by treatment for 10 days with ERKi or WEE1i or the combination. Representative images are shown; scale bar is equivalent to 200 µm (n = 4). *B*, quantification of CellTiter-Glo 3D cell viability assay in organoids at single dose combination from panel *A*. *C*, organoids from (A) were quantified using CellTiter-Glo 3D cell viability assay across a range of WEE1i (adavosertib) and ERKi (SCH772984) doses. Percent viability was normalized to DMSO-treated organoids at the assay endpoint.

## Discussion

It is now well-appreciated that the effectiveness of pharmacologic inhibition of key cancer drivers is offset by the induction of compensatory activities (17,18,56,57). Therefore, identification of treatment-induced compensatory mechanisms can guide the development of combination approaches to better achieve long-term and durable clinical responses. The recent development of direct inhibitors of KRAS (6, 7) has made it even more critical to understand the compensatory activities that can drive resistance to KRAS suppression. In the present study, we applied an activity-dependent chemical proteomics strategy, MIB/MS, to profile the KRAS-dependent kinome in KRAS-mutant PDAC. We identified a diverse spectrum of kinases that were downregulated or upregulated in response to KRAS depletion; many of these had not previously been associated with KRAS effector signaling. We speculated that both groups of kinases may be effective therapeutic targets, with upregulated kinases representing compensatory mechanisms that drive resistance to KRAS suppression. We identified upregulation of DDR1 as one such mechanism and showed that DDR1 inhibition suppressed the growth of KRAS mutant PDAC lines. Conversely, our characterization of one downregulated kinase, WEE1, showed that this DNA damage checkpoint inhibitor can also serve as a therapeutic target. Our study demonstrates the utility of kinome-wide profiling to identify novel strategies for targeting KRAS-mutant PDAC.

Our application of the MIB/MS proteomics screen identified a spectrum of kinases that were down- or upregulated upon acute KRAS suppression. We detected a total of 227 kinases in one or more of the six KRAS-mutant PDAC cell lines analyzed. Of these, 125 (55%) were altered in activation and/or expression in one or more lines upon KRAS suppression, underscoring the diverse consequences that aberrant KRAS function confers upon the human kinome. It is likely that even this is a substantial underestimate: the human kinome encodes over 500 protein kinases. Some kinases were not detected here because they are not recognized by any one of the six broad-spectrum kinase inhibitors used in the affinity purification step; others were not detected due to low or lack of expression in pancreatic cancer (58).

Further, although the genetic landscape of PDAC is dominated by the frequent occurrence of alterations in only four genes (*KRAS*, *TP53*, *CDKN2A* and *SMAD4*), the majority of genetic alterations are found at frequencies of less than 5% (59). Of the 125 KRAS-deregulated kinases, only 28 (22%) were altered across all six KRAS-mutant PDAC lines. Our utilization of additional cell lines for RPPA analysis and follow-up validation further underscored the variability between KRAS-mutant PDAC cell lines. All cell lines used in this study are dependent on mutant KRAS, but have varied dependency on ERK activity (14). This observed heterogeneity is thus consistent with the genetic heterogeneity of PDAC, and highlights the distinct and significant impact that co-occurring mutations may have on KRAS function, and, consequently, on response to specific therapies.

We observed significant overlap of the kinase activity/expression changes caused by acute genetic suppression of KRAS compared with those caused by pharmacologic inhibition of ERK. Nine of 13 KRAS-downregulated kinases and four of 15 KRAS-upregulated kinases were also altered upon ERK inhibition. That we observed a dominant role of the ERK MAPK cascade in regulating the KRAS-dependent kinome is consistent with the critical role of this protein kinase cascade in driving KRAS-dependent PDAC growth. Nevertheless, many of the 28 KRAS-regulated kinases are not known to be directly associated with ERK signaling. Only eight (CKIIε, CDK1, CHK1, ERK1, EPHA2, SRPK2, TNIK and TTK) are listed in a recent compilation of direct or indirect ERK substrates from 14 different phosphoproteomic studies (11). However, it is still possible that some of the other 20 KRAS-regulated kinases are ERK substrates. The compilation includes no studies performed in PDAC. In our ongoing phosphoproteomic analyses of ERK-dependent phosphorylation in PDAC, we have identified numerous potential direct or indirect ERK substrates that have not been identified previously. Finally, none of the 28 kinases identified in the present study are listed in the GSEA KRAS gene signature, indicating that many of the kinases we detected are altered in activity rather than by gene expression.

Finally, some observed alterations may be indirect rather than associated specifically with KRAS signaling. For example, reduced kinase expression may be a consequence of the incomplete G1 arrest caused by KRAS suppression. NEK2 is undetectable during G1, but accumulates progressively throughout S phase, reaching maximal levels in late G2 (60).

Our working hypothesis for this study was that the kinases that are upregulated upon KRAS suppression would be distinct from those typically utilized by KRAS for its transforming activities. Instead, it is now well-established that compensatory kinome reprogramming occurs upon pharmacologic inhibition of RAF and PI3K signaling (23, 24). We hypothesized that loss of KRAS signaling would similarly induce compensatory kinome reprogramming, such that the upregulated kinases would enable maintenance of the transformed phenotype. Consistent with this possibility was the significant enrichment of upregulated tyrosine kinases, a class that includes many oncogenic kinases (e.g., EGFR, HER2, etc.). To evaluate this possibility, we selected two kinases, JAK1 and DDR1, for validation analyses. Since there is evidence that JAK1 can act as a cancer driver in multiple cancer types (61), we were surprised that pharmacologic inhibition of JAK1, either alone or together with ERK inhibition, did not negatively impact PDAC growth. This finding emphasizes that the potential therapeutic value of an upregulated or hyperactivated kinase requires functional validation, not simply evidence of increased expression or activity.

In contrast to JAK1, we verified that pharmacological inhibition of DDR1 alone or in combination with an ERK inhibitor caused additive or synergistic growth suppression of PDAC cell lines. To date, there has been limited effort in the pursuit of DDR1 as a therapeutic target in PDAC and there are no clinically tractable DDR1-specific inhibitors. However, the DDR1- selective pharmacological inhibitor 7rh, together with chemotherapy, impaired tumorigenic growth of PDAC cell lines (44). In a complementary study, genetic ablation of *Ddr1* in the KPC (*Kras*^G12D/+^; *Tp53*^R172/+^) mouse model of PDAC impaired metastatic tumor growth (43). Finally, a recent kinome profiling screen using a different experimental method than MIB/MS demonstrated upregulation of DDR1 expression in patient-derived PDAC cell lines (62). In agreement, we observed that direct pharmacologic inhibition of DDR1 with the small molecule inhibitor 7rh suppressed PDAC cell proliferation in vitro. Additionally, we observed synergistic reduction in proliferation when we combined the DDR1 inhibitor with the ERK1/2-selective inhibitor SCH772984, supporting further evaluation of this combinatorial therapeutic strategy.

The downregulated kinases were enriched (11 of 13) in kinases that modulate cell cycle progression, particularly processes involved in mitosis. Inhibition of some of these individual kinases are sufficient to impair cancer growth, indicating that KRAS regulates growth through a multitude of mechanisms. For example, polo-like kinase (PLK1) has been identified as a synthetic lethal interactor with mutant RAS (63). We showed previously that pharmacologic inhibition of TTK impaired PDAC growth (64). In agreement with previous studies (59,65–67), we showed here that inhibition of WEE1 suppresses PDAC growth. Additionally, we found that concurrent ERK inhibition, to counter the compensatory ERK activation associated with WEE1 inhibition, further enhanced WEE1i growth-inhibitory activity. Strikingly, concurrent WEE1i and ERKi effectively reduced growth of both 2D adherent PDAC cultures as well as 3D organoid cultures. The WEE1i adavosertib is currently being evaluated alone or in combination with other anti-cancer therapies in 20 active or recruiting clinical trials, including one in pancreatic cancer (NCT02194829, accessed Feb. 22, 2021).

In summary, the MIB/MS chemical proteomics strategy provides a powerful experimental approach to delineate a more complete understanding of the KRAS-dependent kinome, implicating kinases not identified by other methods. We additionally showed that both downregulated and upregulated kinases upon loss of KRAS represent potential therapeutic targets for PDAC treatment. Our findings also further establish mechanisms by which the ERK MAPK effector pathway drives KRAS-dependent PDAC growth, through regulation of multiple distinct regulators of cell cycle progression, particularly mitosis. Finally, that KRAS suppression caused upregulation of a diverse spectrum of functionally distinct kinases underscores the need to develop combination inhibitor therapies to thwart treatment-induced kinome reprogramming, that will result in acquired resistance and limit the long-term efficacy of mutant-specific KRAS inhibitors.

## Experimental procedures

### Cell culture

Cell lines were obtained from the American Type Culture Collection (AsPC-1, Panc10.05, SW-1990, MIA PaCa-2, PANC-1, HPAC and HPAF-II) or were a gift from J. Fleming at MD Anderson Cancer Center (Pa01C, Pa02C, Pa14C and Pa16C). Cells were maintained in either Dulbecco’s minimum essential medium (DMEM) (Gibco, 11995-065; MIA PaCa-2, PANC-1, HPAC, HPAF-II, Pa01C, Pa02C, Pa14C and Pa16C) or RPMI-1640 (Gibco; AsPC-1, Panc 10.05 and SW-1990) supplemented with 10% fetal bovine serum (Sigma-Aldrich), penicillin and streptomycin (Sigma-Aldrich). All cell lines were short-tandem repeat (STR)-profiled to confirm their identity. All cell lines tested negative for mycoplasma contamination.

### Antibodies and reagents

SCH772984 was provided by Merck. The following compounds were purchased from Selleckchem: adavosertib (S1525), filgotinib (S7605), ruxolitinib (S5243), and tofacitinib (S2789); or from Sigma-Aldrich: 7rh (SML1832), MG-132 (M7449). The following antibodies were obtained from Cell Signaling Technology: anti-DDR1 (5583S), anti-PYK2 (3090S), anti-STAT3 (9139S), anti-phosphorylated STAT3 Tyr^705^ (9145S), anti-WEE1 (13084S), anti-CHK1 (2360S), anti-PLK1 (4513T), anti-ERK (4696S), anti-phosphorylated ERK Thr^202^/Tyr^204^ (4370S), anti-phosphorylated CDC2 Tyr^15^ (4539S), anti-phosphorylated HistoneH3 Ser^10^ (53348S); from Sigma-Aldrich: anti-vinculin (V9131), anti-β-actin (A5441) and anti-KRAS (WH0003845M1); from Invitrogen: anti-phosphorylated PYK2 Tyr^402^ (44-618G); or from Santa Cruz Biotechnology: anti-CDC2 (sc-54).

### RNAi knockdown studies

Short-interfering RNA (siRNA) experiments were performed with 10 nM siRNA and RNAiMAX Lipofectamine (Invitrogen, 13778150) according to the manufacturer’s protocol. siRNAs were obtained from Thermo Fisher: Non-specific siRNA (4390844), siKRAS-1 (4390825-s7939), siKRAS-2 (4390825-s7940), siWEE1-1 (4390824-s21), siWEE1-2 (AM51331-404), siDDR1-1 (4390824-s2298), and siDDR1-2 (4390824-s2230). On day 1, cells were plated at 2 x 10^5^ cells per well of a 6-well plate. On day 0, Lipofectamine was warmed to room temperature and added to Opti-MEM (Gibco, 31985-070) to a final concentration of 20x before low-speed vortex. siRNAs were added to individual tubes of Opti-MEM at 20x concentration. Both Lipofectamine and siRNA tubes were incubated at RT separately for 5 min. After incubation, Lipofectamine in Opti-MEM was combined at a 1:1 ratio with individual siRNAs in Opti-MEM. Tubes were carefully inverted five to seven times to mix Lipofectamine and siRNA in Opti-MEM and were subsequently incubated at RT for 30 min. Two hundred µl of the Lipofectamine with siRNA were added to 1.8 ml of fresh cell culture media per well. Cells were incubated for specified times before collection for immunoblotting, proliferation, clonogenic, cell cycle or apoptosis assays.

For qRT-PCR experiments, the above transfection protocol was modified to combine the day 1 and day 0 steps. Cells (2 x 10^5^) were plated in 1.8 ml cell culture medium with the addition of 200 µl of Lipofectamine with respective siRNA on day 0.

### Multiplexed inhibitor beads-mass spectrometry (MIB/MS)

Samples were prepared according to the RNAi knockdown protocol outlined in Experimental proceedures. Following 72 h of RNAi knockdown with either non-specific siRNA or siKRAS 1, the cell plates were placed on ice and the cells were washed 5x with large-volume, ice cold phosphate-buffered saline (PBS). The final PBS wash was thoroughly aspirated and MIB/MS lysis buffer (50 mM HEPES (pH 7.5), 0.5% Triton X-100, 150 mM NaCl, 1 mM EDTA, 1 mM EGTA, 10 mM sodium fluoride, 2.5 mM sodium orthovanadate, protease inhibitor cocktail (Roche), 1% phosphatase inhibitor cocktail 2 (Sigma-Aldrich), and 1% of phosphatase inhibitor cocktail 3 (Sigma-Aldrich) was added to the adherent cells and permitted to incubate on ice for 5 min. Using a rubber cell scraper, cell lysates were transferred from the plates to 1.7 ml microcentrifuge tubes and placed on ice for 10 min. Cell lysates were subsequently sonicated four times for 15 sec at 50% power, alternating between samples to allow cooling on ice between pulses. Cell lysates were then centrifuged at top speed (13,200 rpm) for 10 min at 4°C. Supernatants were clarified using a 0.2 µm filter, transferred into labeled tubes and stored at -80°C (∼5 mg protein per experiment). A small portion of lysate was removed for Bradford assay estimation of protein concentration. The lysates were thawed and gravity-flowed over multiplexed kinase inhibitor beads (MIBs; Sepharose conjugated to VI-16832, CTx-0294885, PP58, Purvalanol B, UNC8088A, and UNC21474). MIBs were then washed with high salt (1 M NaCl) and low salt (150 mM NaCl + 0.1% SDS) lysis buffers without the inhibitors. The samples were boiled with the elution buffer (100 mM tris-HCl, 0.5% SDS, and 1% β-mercaptoethanol, pH 6.8) at 100°C for 5 min to elute the bound kinases from MIBs. The eluted kinases (proteins) were concentrated with Amicon Ultra-4 (10K cutoff) spin columns (Millipore), purified by removing the detergent using methanol/chloroform extraction, and digested by sequencing grade trypsin (Promega) overnight at 37°C. Hydrated ethyl acetate extraction was used to remove Triton, and PepClean C18 spin columns (Thermo Scientific) were used to de-salt the digested peptides.

Biological triplicates of the MIB samples were analyzed by LC-MS/MS as we described previously (33). Briefly, each sample was injected onto an EASY-Spray PepMap C18 column (75 mm id 3 25 cm, 2 mm particle size) (Thermo Scientific) and separated over a 2-h method. The gradient for separation consisted of 5%–32% mobile phase B at a 250 nl/min flow rate, where mobile phase A was 0.1% formic acid in water and mobile phase B consisted of 0.1% formic acid in ACN. The Thermo QExactive HF was operated in data-dependent mode where the 15 most intense precursors were selected for subsequent HCD fragmentation (set to 27%).

Following MaxQuant processing of data and generation of Label-free quantification (LFQ) intensity values, data files were analyzed using R (version 3.5.2). Kinases were excluded if they were not present in >50% of samples or if they contained two or more peptides. For imputation of missing values, a normal distribution was modeled on the non-missing LFQ intensity values of the kinases containing missing intensity values. Imputed values were drawn randomly from this distribution. Following filtering and imputation, LFQ intensity values were log2 transformed and the fold change over the median vehicle value was calculated for each kinase. Kinases significantly different between treatments with siNS and siKRAS, or with DMSO and ERKi, were determined using one-way ANOVA (Benjamini-Hochberg adjusted p value < 0.05). Euclidean distance and average linkage were utilized for unsupervised hierarchical clustering of log2 fold-change values for significant kinases.

### Immunoblotting

Plates containing the PDAC cell lines were washed with PBS and lysed with Triton X-100 lysis buffer (25 mM tris buffer, 100 mM NaCl, 1 mM EDTA and 1% Triton X-100), supplemented with cOmplete protease inhibitor (Roche) and Phosphatase Inhibitor Cocktail (Sigma-Aldrich). Cells were incubated on ice for 10 min before scraping into a microcentrifuge tube and placed back on ice. Sample tubes were centrifuged for 10 min at 13,200 rpm at 4°C. The supernatant was collected and used to determine protein concentration using a Bradford Assay (Bio-Rad), and samples were prepared with 4x Laemmli Sample Buffer (Bio-Rad). Samples were then boiled and stored at -20°C. Ten percent polyacrylamide gels were used to clarify lysates before transferring onto PVDF membranes (Millipore, IPVH00010) at 90 V for 90 min. Membranes were blocked for 1 h in 5% BSA solution diluted in TBST (TBS with 0.05% Tween-20) and washed with TBST. Membranes incubated overnight in primary antibodies diluted in 5% BSA solution supplemented with sodium azide. Secondary antibodies used were goat anti-mouse (Invitrogen, 31432) and goat anti-rabbit (Invitrogen, 31462). Membranes were washed with TBST before imaging using the ChemiDoc MP Imaging System (Bio-Rad) and ECL reagent.

### Quantitative real-time PCR

Cells were grown in 6-well plates, transfected with siRNA as indicated, and harvested. RNA was extracted using the RNeasy Isolation Kit (Qiagen) and converted to cDNA using the High-Capacity cDNA Reverse Transcription Kit (Applied Biosystems). RT-PCR was performed using the TaqMan system (Applied Biosystems) in a 384-well format. FAM-labeled target primer and endogenous human ACTB control (beta actin) (Thermo Fisher Scientific) were mixed with master mix and template, and after 40 cycles were analyzed on an QuantStudio 6 Flex Real-Time PCR System (Applied Biosystems).

### Flow cytometry

For apoptosis assays, TACS Annexin V-FITC Kits (BD Biosciences) were utilized to measure apoptosis. The following protocol is closely adapted from the manufacturer’s specifications. Prior to trysinization, floating cells in the spent culture medium were collected. Cells were then trypsinized cells and collected, mixed, and centrifuged at 300*g* for 5 min at room temperature. The cells were washed with phosphate buffered saline (PBS) and centrifuged at 300*g* for 5 min before incubating the cell pellet in annexin V staining solution (1% Annexin V-FITC, 1x propidium iodide solution, in 1x calcium-containing binding buffer) in the dark for 15 min at room temperature. Cell mixture was subsequently diluted 1:5 in binding buffer. For cell cycle analysis, adherent cells were washed with PBS prior to being trypsinized. After trypsinization, cells were centrifuged at 300*g* for 5 min before washing once in PBS. Cells were pelleted and resuspended in 1 ml PBS before adding 9 ml of 70% ethanol drop-wise to each tube with gentle agitation. Cells were permitted to fix for a minimum of 18 h at 4°C. Fixed cells were then pelleted, washed once in PBS, resuspended in 40 μg/ml propidium iodide (PI), 100 μg/ml RNase A in PBS, and incubated at 37°C for 3 h.

Apoptosis assays were analyzed using FCS Express. A FSC-A vs. SSC-A dot plot was used to exclude debris and generate a “cells” gate. “Cells” were plotted in a FITC-A (x) vs. PI-A (y) dot plot and apoptotic cells (FITC+) were analyzed. Cell cycle analyses were performed with FCS Express. After first establishing a “cells” gate, a “singlets” gate was determined using a FSC-A (x) vs. FSC-H (y) dot plot. Singlets were then analyzed in a histogram for PI-A content before employing the Multicycle algorithm to analyze cell cycle.

### Reverse Phase Protein Array (RPPA)

Samples were prepared according to the RNAi knockdown protocol outlined in Experimental procedures. Following 72 h of RNAi-mediated knockdown with non-specific siRNA, siKRAS-1, or siKRAS-2, plates were washed three times with PBS. Cell lysates were prepared as previously described (28). Coomassie Protein Assay Reagent kit (Thermo Fisher Scientific) was used to measure protein concentration, following the manufacturer’s instructions. Tris-glycine SDS 2x sample buffer (Life Technologies) with 5% β-mercaptoethanol was used to dilute cell lysates to 0.5 mg/ml. Samples were boiled for 8 min and stored at -20°C until arrayed. An Aushon 2470 automated system (Aushon BioSystems) (68) was used to immobilize cell lysates and the internal controls and print in technical replicates (n = 3) onto nitrocellulose-coated glass slides (Grace Bio-Labs). Sypro Ruby Protein Blot Stain (Molecular Probes) was used to quantify protein concentration in each sample, following the manufacturer’s instructions. Reblot Antibody Stripping solution (Chemicon) was used to pretreat the remaining arrays (15 min at room temperature). The arrays were washed with PBS and incubated in Iblock (Tropix) for 5 h before antibody staining (34). Arrays were incubated with 3% H_2_O_2_, avidin, biotin (DakoCytomation), and an additional serum-free protein block (DakoCytomation) to reduce nonspecific binding of endogenous proteins. Staining was performed using an automated system (DakoCytomation) was used. Each slide was probed for 30 min with one antibody targeting the protein of interest, with 149 antibodies that target proteins involved in signaling networks that regulate cell growth, survival and metabolism were used to probe arrays (table S3). All antibodies used were validated as described previously (69). Signal amplification was determined by using biotinylated anti-rabbit (Vector Laboratories) or anti-mouse secondary antibody (DakoCytomation) and a commercially available tyramide-based avidin-biotin amplification system (Catalyzed Signal Amplification System, DakoCytomation). IRDye 680RD streptavidin (LI-COR Biosciences) fluorescent detection system was used. TECAN laser scanner was used to scan Sypro Ruby and antibody-stained slides, and the images were analyzed using commercially available software (MicroVigene Version 5.1.0.0, Vigenetech) as previously described (70).

### Proliferation assays

For 96-well-format growth assays, 10^3^ cells per well were plated on Day 1 in 200 µl cell culture media. Following overnight incubation, one plate per cell line was isolated on Day 0 and quantified by counting calcein AM positive (500 nM, 20 min) labeled live cells using the SpectraMax i3x multimode detection platform (Molecular Devices) for the assessment of plating efficiency. Next, we utilized the TECAN 300D dispenser to add inhibitor titrations to the plates. Dose ranges used are indicated in figures and/or figure panels. DMSO was kept at or below 0.01% final concentration. Cells were incubated at 37°C and 5% CO_2_ for 5 days. Plates were quantified using calcein AM staining (500 nM, 20 min) followed by counting of positively stained cells using the SpectraMax i3x multimode detection platform. Raw cell numbers were minimally adjusted by cell line depending on the plating efficiency determined on Day 0. Next, raw cell counts (positively labeled with calcein AM) were normalized to the untreated wells. One hundred % growth was assigned to an average of the untreated well cell counts, all treated wells were calculated relative to this number and technical replicates were averaged together. Growth curves were constructed by first modeling the raw data with the drc R package (version 3.0-1) and the four-parameter log-logistic function LL.4 (4PL). Next, we isolated the GI_50_ values from the models and drew the growth curves using the ggplot2 package in R (version 3.3.2).

### BLISS synergy analysis

First, quantifications of proliferation assays (2D or 3D cultures) were normalized to untreated controls (100% growth). Next, treatment effect sizes were calculated by subtracting the growth percentage (as a decimal) from 1. Values above 1 were removed from BLISS analysis (corresponding to treatments that stimulated cell growth above untreated control). Expected effect sizes for each treatment combination were calculated according to the BLISS algorithm (71). Expected effect size was then divided by observed effect size and the results were mapped to a color scale (0-1 are shades of red, indicative of “synergy”; 1 is white, indicative of additivity; greater than 1 are shades of blue, indicative of “antagonism”) and plotted as a heatmap using the ComplexHeatmap package in R (version 2.4.3).

### Proteasome inhibitor studies

Knockdown was performed in the manner described in Experimental procedures using either non-specific siRNA, siKRAS 1, or siKRAS 2. Following incubation with respective siRNA for 68 h, the cell culture medium was replaced with standard cell culture medium supplemented with vehicle or 10 µM MG132. The cells were then incubated at 37°C for 4 h before using the standard immunoblotting protocol as described in Experimental procedures.

### Data and materials availability

Data from this study are available in the Supporting information.

## Supporting Information

Data File S1. MIB/MS Log_2_-LFQ values following 72 h siNS or siKRAS1

Data File S2. Reverse Phase Protein Array following 72 h siNS or siKRAS

Data File S3. MIB/MS Log_2_-LFQ values following 24 h DMSO or SCH772984 (ERKi)

## Acknowledgements

We thank Sarah Howard for her assistance in manuscript preparation.

## Author contributions

J.N.D. and C.J.D. designed the experiments. J.N.D., K.R.S., P.S.H., A.D.C. and C.J.D. wrote the manuscript. J.N.D., J.E.K., K.R.S., D.R.B., P.S.H., Z.D.K., B.P., R.Y., and R.G.H. performed experiments. T.S.K.G., M.P., E.B., L.E.H., and E.F.P. provided proteomics analyses; J.N.D. performed bioinformatics analyses; N.U.R., L.E.H., L.M.G., A.D.C., and C.J.D. provided scientific guidance.

## Abbreviations

2D: two-dimensional
3D: three-dimensional
CRISPR: clustered regularly interspaced short palindromic repeats
DDR1: discoidin domain receptor 1
ERKi: ERK inhibition
FBS: fetal bovine serum
GTP: guanosine triphosphate
GI_50_: concentration at which growth is 50% of maximum
IC_50_: concentration at which the target is 50% inhibited
MAPK: mitogen-activated protein kinase
MEKi: MEK inhibition
MIB/MS: multiplex kinase inhibitor beads/mass spectroscopy
NS: non-specific
PDAC: pancreatic ductal adenocarcinoma
RPPA: reverse phase protein array
RNAi: RNA interference
RTK: receptor tyrosine kinase
siRNA: small interfering RNA
STRING: Biological database of protein-protein interactions
WT: wild-type

---

**Figure S1.**
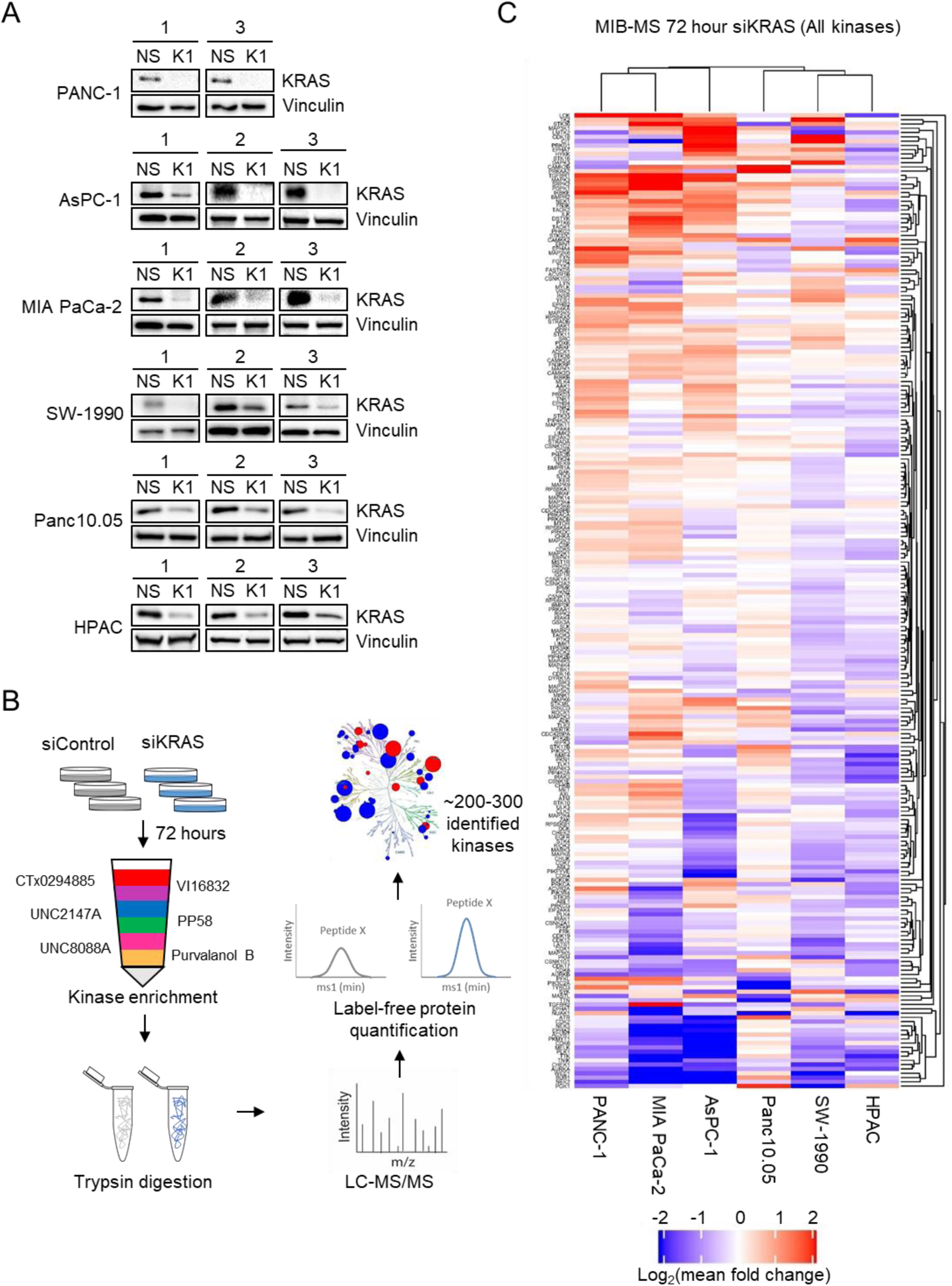
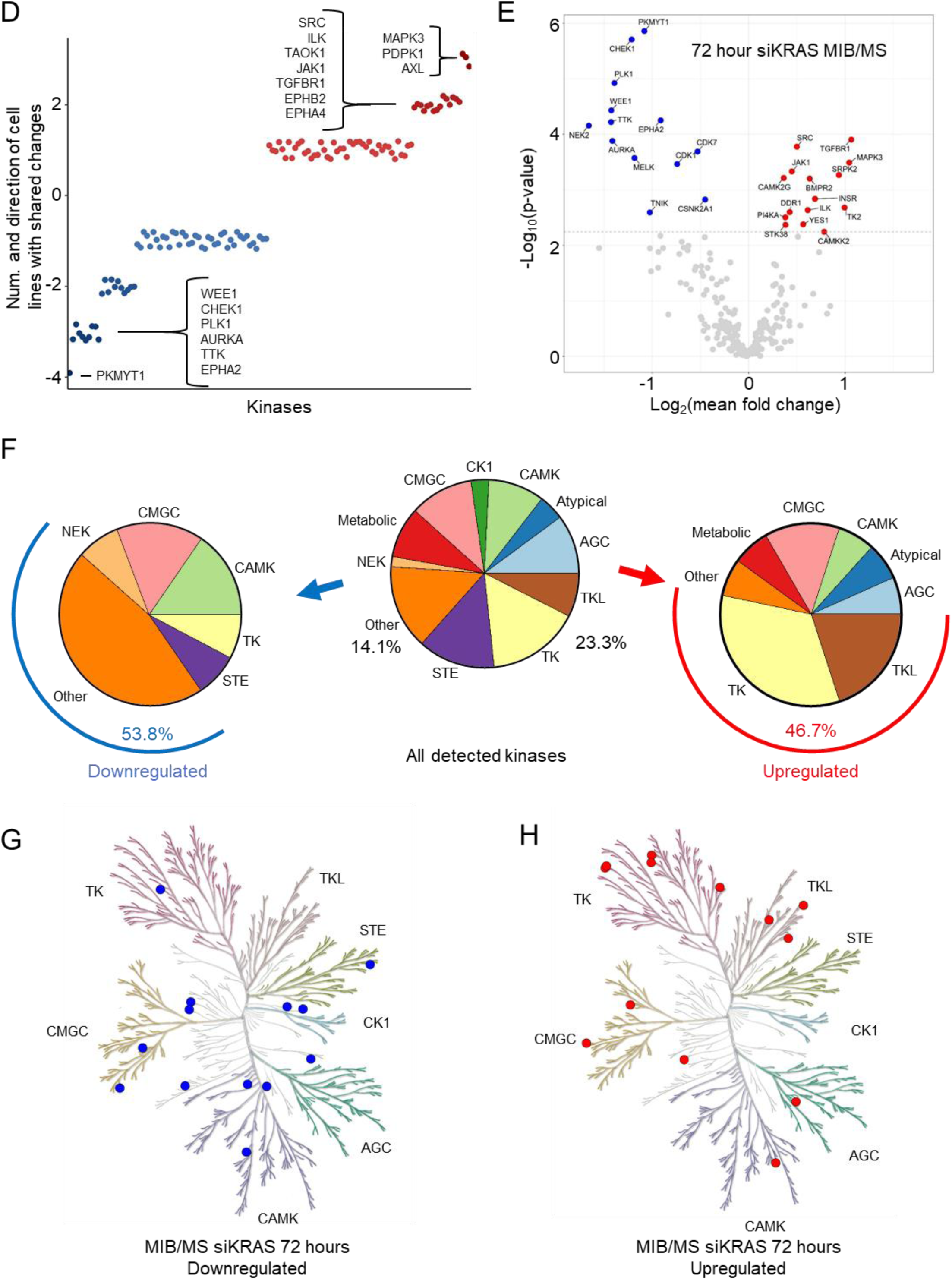
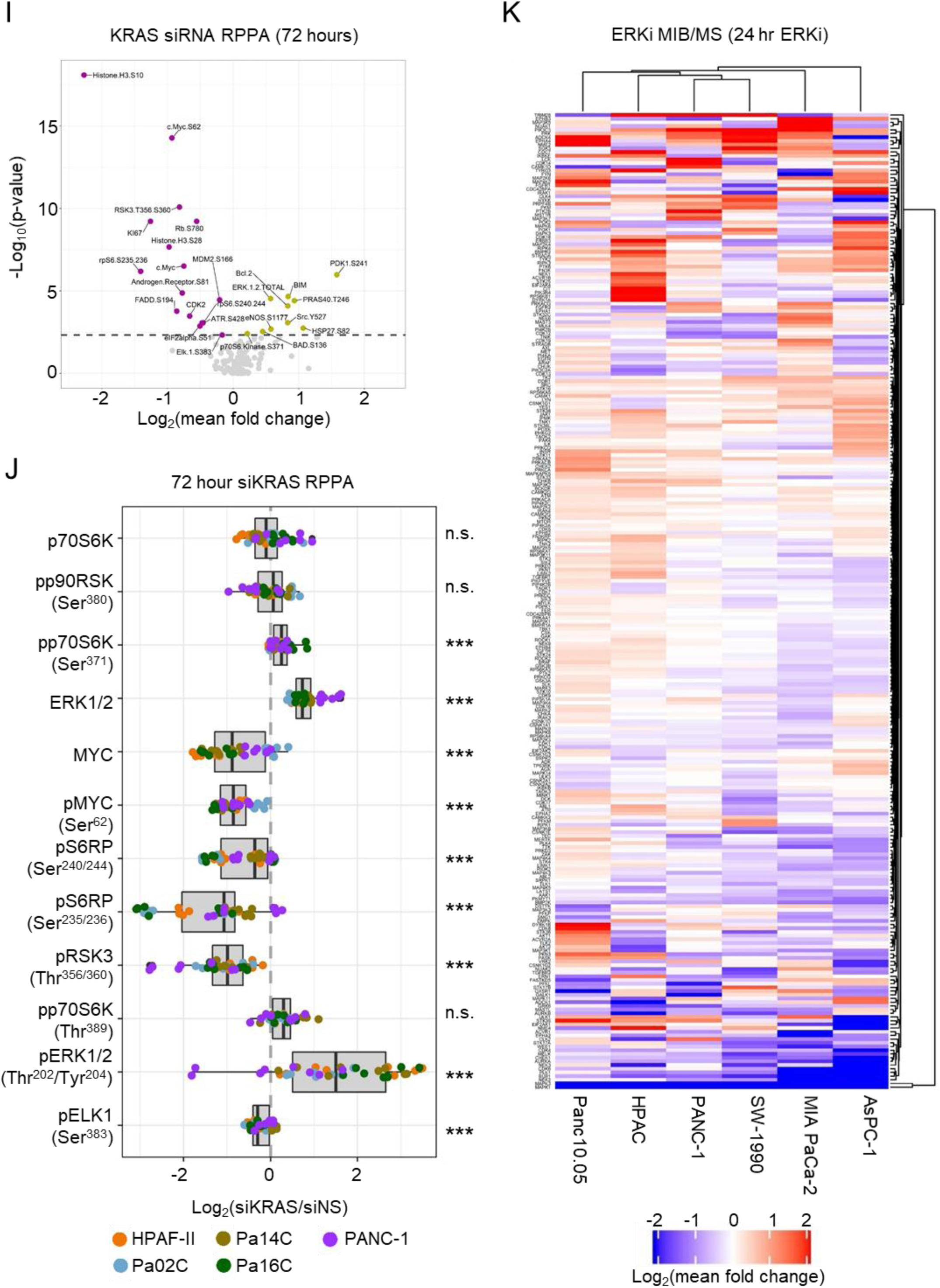
KRAS-dependent kinome-wide alterations. *A,* immunoblot analysis of siNS- and siKRAS-treated samples submitted for MIB/MS analysis. Prior to submitting sample lysates to the UNC Proteomics Core, a small portion of the lysate was retained to confirm KRAS knockdown. Replicate 2 of PANC-1 was lost during processing. *B,* schematic of MIB/MS method performed. Cells were transfected with respective siRNA oligonucleotides for 72 h prior to cell lysis and kinase enrichment. *C,* heatmap summarizing all 227 kinases detected among six KRAS mutant, KRAS-dependent PDAC cell lines (SW-1990, MIA PaCa-2, PANC-1, HPAC, AsPC-1 and Panc10.05). Log_2_(fold-change) values were calculated by comparing siKRAS and siNS for each cell line. *D,* kinase activity/expression changes identified from siKRAS MIB/MS (Fig. S1*C*) in individual cell lines were compared. All 125 kinases significantly (p adjusted < 0.05) altered in at least one cell line are displayed here. *E,* volcano plot showing results of significance testing after ANOVA [y-axis] with the log-transformed mean fold change in siKRAS over siNS (72 h). Significant kinases are colored red (increased) or blue (decreased) if p-value was significant following Benjamini-Hochberg correction (n = 3). *F,* pie charts indicating the proportion of kinase class representation in all detected kinases (227), upregulated kinases (15), and downregulated kinases (13). *G-H,* Tree diagrams generated with KinMap (74) showing kinase family relationships of upregulated (J) or downregulated (K) kinases detected in 72 h siKRAS samples from Fig 1A. *I,* volcano plot highlighting significantly altered proteins/phosphoproteins identified by RPPA analysis (p adjusted < 0.05). Five PDAC cell lines (HPAF-II, Pa02C, Pa14C, Pa16C and PANC-1) were transfected with siNS or siKRAS for 72 h prior to sample collection. Significant kinases are colored gold [increased] or violet [decreased]. Data are representative of five independent biological replicates. *J,* selected results from RPPA analysis are shown for samples from *I*. ANOVA was used to determine significant alterations among all cell lines (n.s. indicates non-significant; * p value < 0.05; ** p value < 0.005; *** p value < 0.0005). *K,* heatmap summarizing all kinases detected among six PDAC cell lines in response to ERK inhibition or DMSO treatment. Values were calculated by comparing ERKi treated samples to DMSO treated samples within each cell line group.

**Figure S2.**
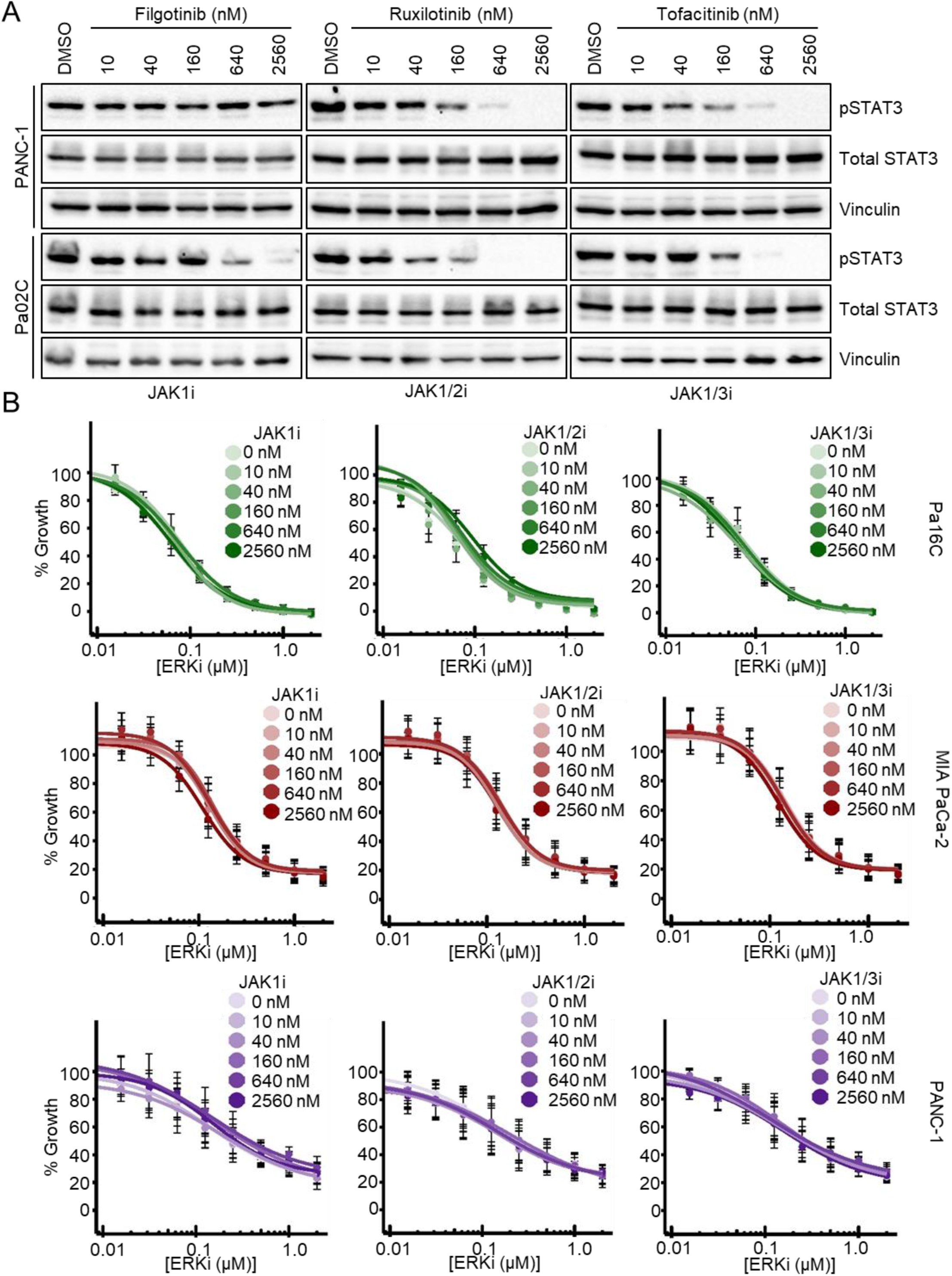
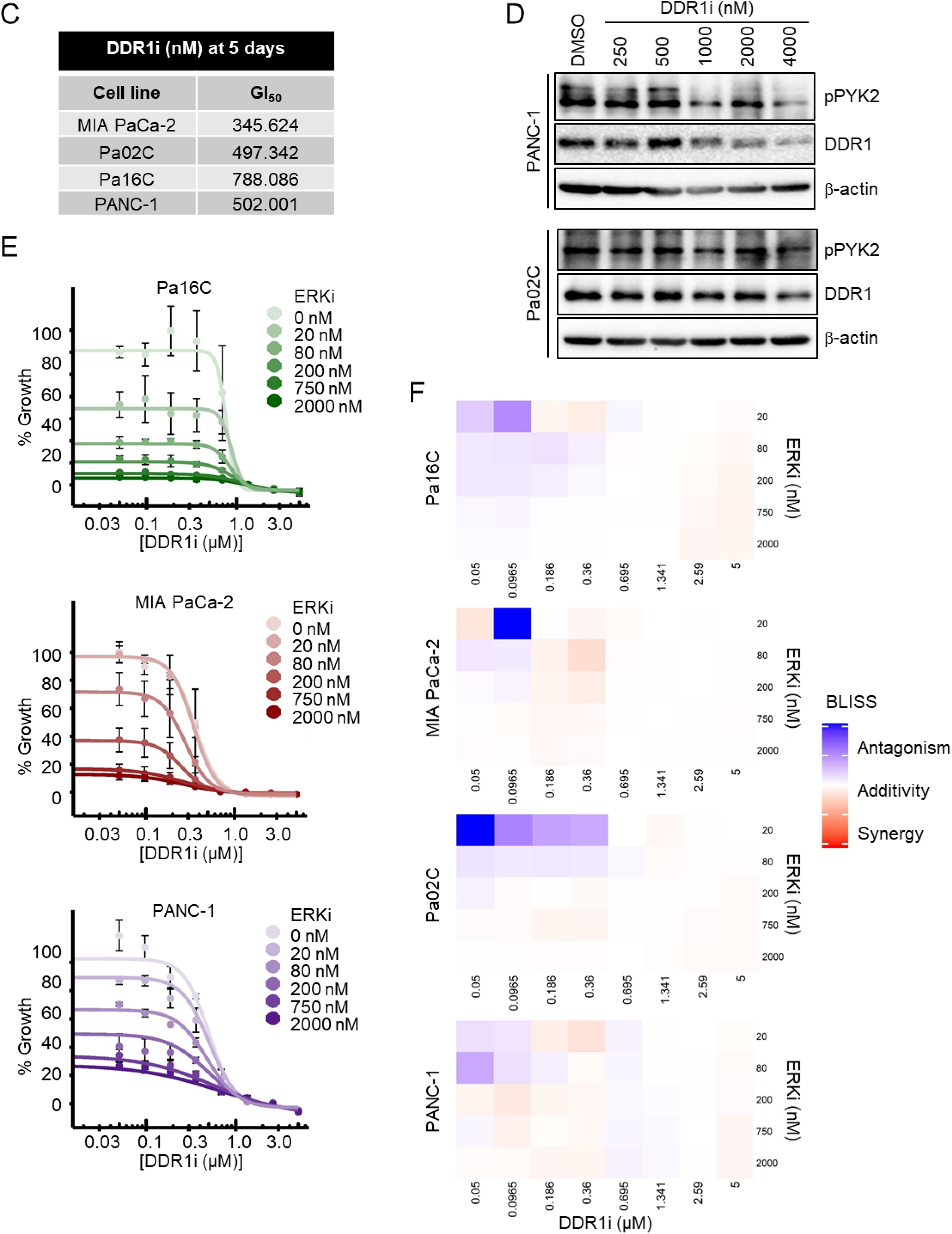
Upregulation of DDR1, but not JAK, contributes to PDAC survival upon KRAS knockdown. *A,* immunoblot analysis of two PDAC cell lines treated for 24 h with JAK1i (filgotinib), JAK1/2i (ruxolitinib), or JAK1/3i (tofacitinib) at increasing concentrations. Data are representative of three independent experiments. *B,* proliferation of PDAC cells following treatment for 5 days with JAK1i (filgotinib), JAK1/2i (ruxolitinib), or JAK1/3i (tofacitinib) in combination with ERKi (SCH772984) at various concentrations. Cell numbers at endpoint were normalized to vehicle-treated control (100% growth) for each cell line. Curves were fit using a 4-parameter log-logistic function and the drm package in R (n=3). *C,* GI_50_ values for the DDR1 inhibitor 7rh (5 days of treatment) were calculated in four PDAC cell lines (n = 3). *D,* immunoblot analysis of PDAC cell lines treated for 24 h with increasing doses of the DDR1 inhibitor 7rh (n = 3). *E,* two-dimensional growth of PDAC cells following treatment for 5 days with DDR1i (7rh) and ERKi at various concentrations in combination. Quantification and analysis is the same as in panel *A* (n = 3). *F,* BLISS Synergy scores were calculated using the proliferation effect sizes from Fig. 2D. Scores < 1 indicate synergy (red), scores = 1 are additive (white), and scores > 1 are antagonistic (blue) (n = 3).

**Figure S3.**
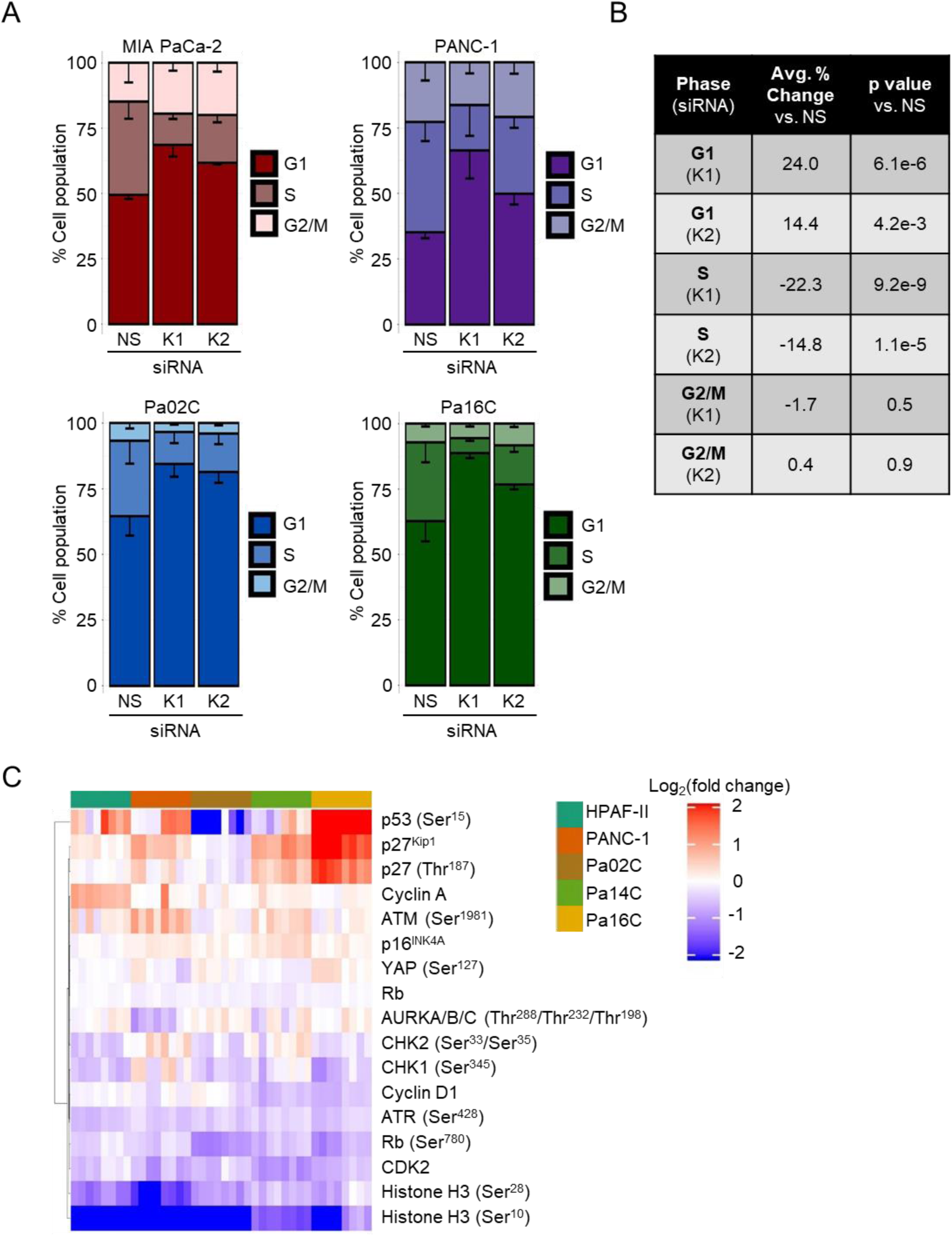
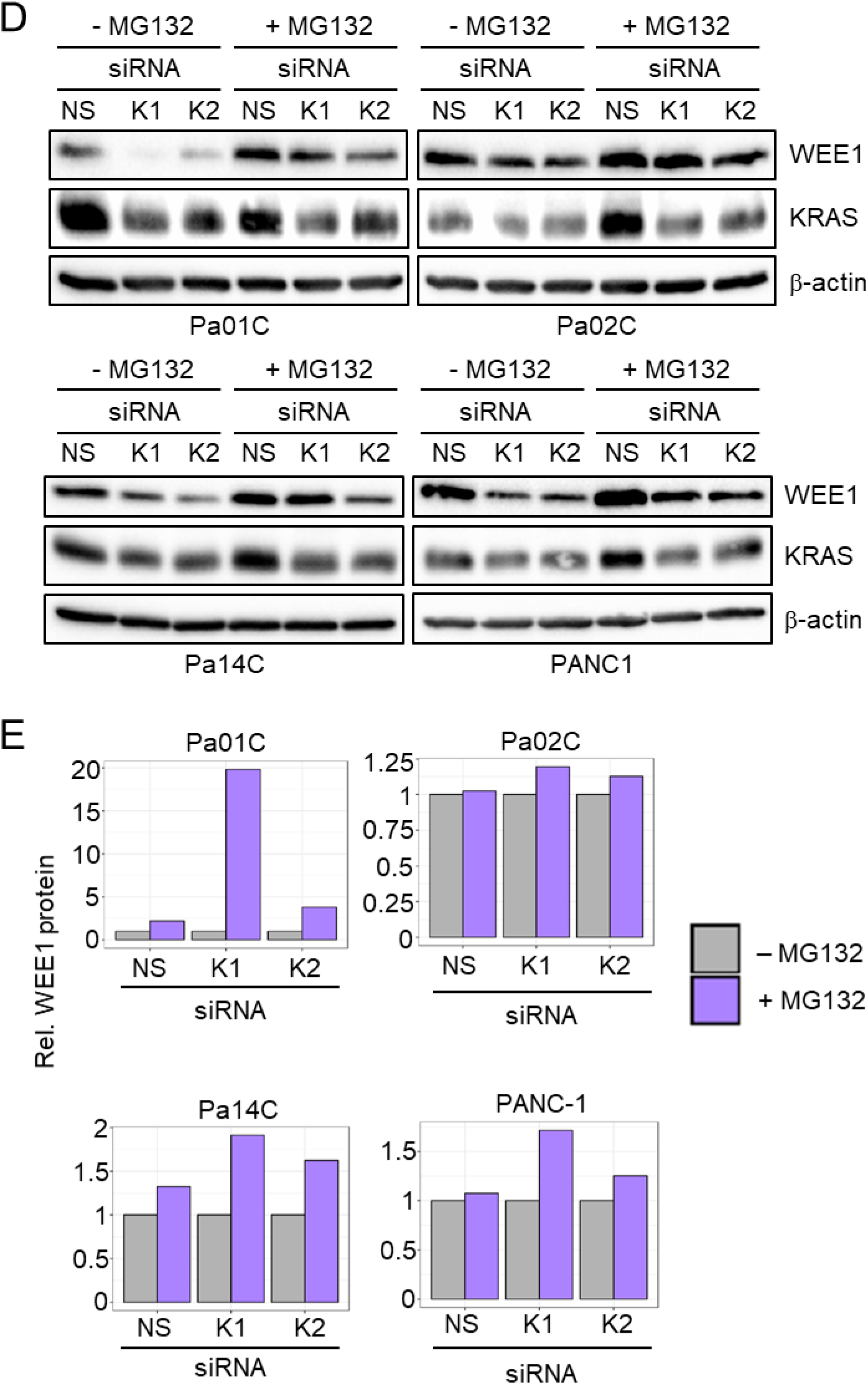
KRAS or ERK suppression causes loss of checkpoint and DNA damage response kinases. *A,* flow cytometry cell cycle analysis was performed following 72-h siNS, siKRAS 1 and siKRAS 2 treatment of four PDAC cell lines. Quantification was performed in FCS Express (n = 3). *B,* tabular summary of results from panel *A*. Student’s *t*-test was used to identify significant differences between KRAS knockdown and control. *C,* reverse phase protein array (RPPA) analysis of 5 PDAC cell lines following 72-h siRNA knockdown of KRAS. Relative color of heatmap indicates the log2-transformed fold change of siKRAS over siNS, with red indicating a greater value in siKRAS compared to siNS and blue indicating a lesser value in siKRAS compared to siNS. All 4 replicates for both siKRAS constructs (K1, K2) are shown as individual columns (n = 4). *D,* protein stability of WEE1 as determined by 4-h MG132 proteasomal inhibitor treatment following 26-h treatment with non-specific (NS) or KRAS-targeted (K1, K2) siRNA (n = 3). *E,* quantification of blot analysis from panel *D*. Densitometry estimates were first normalized for loading efficiency using β-actin. Next, each sample was calculated as a ratio of MG132+/MG132- and paired by siRNA construct.

**Figure S4.**
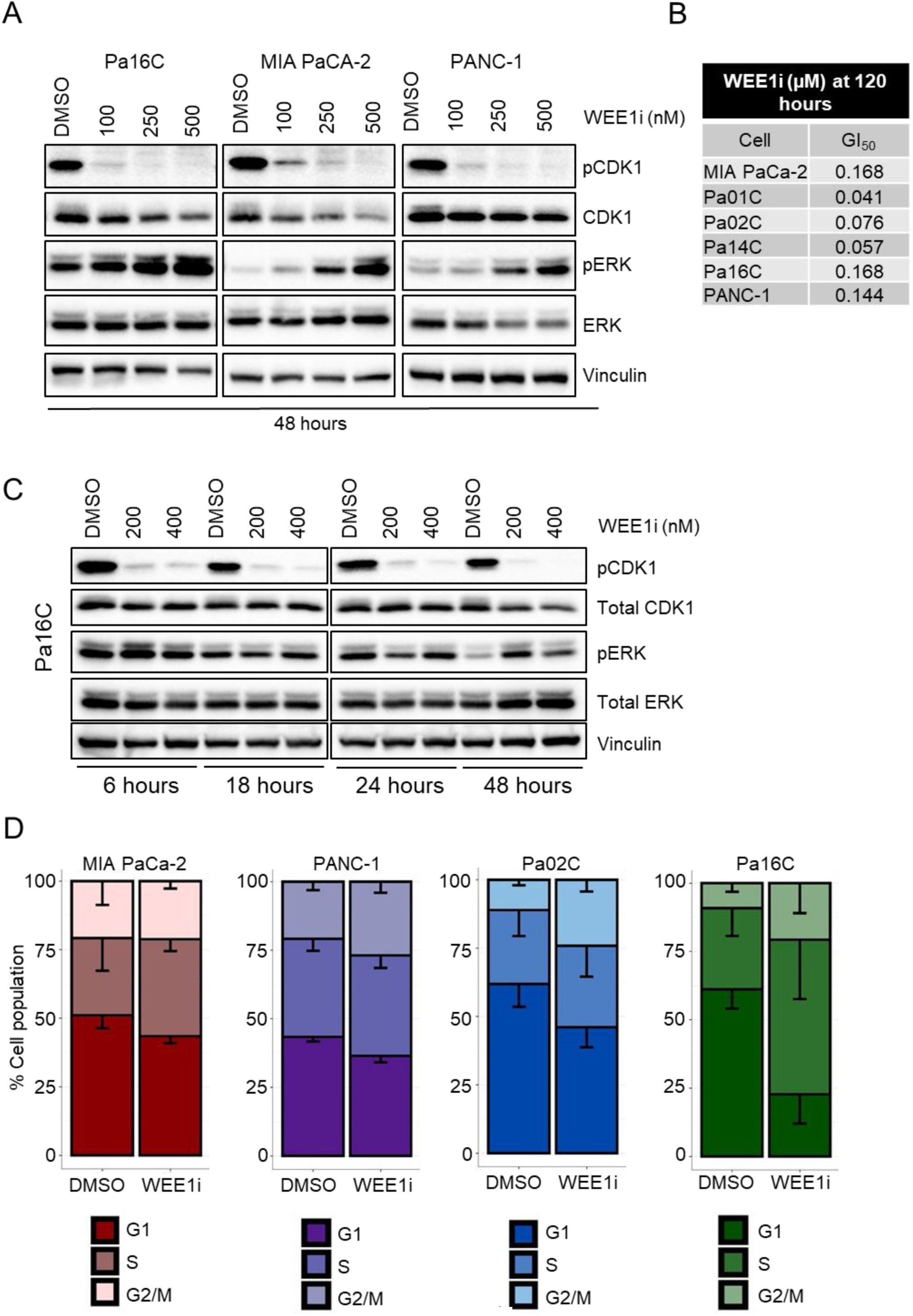
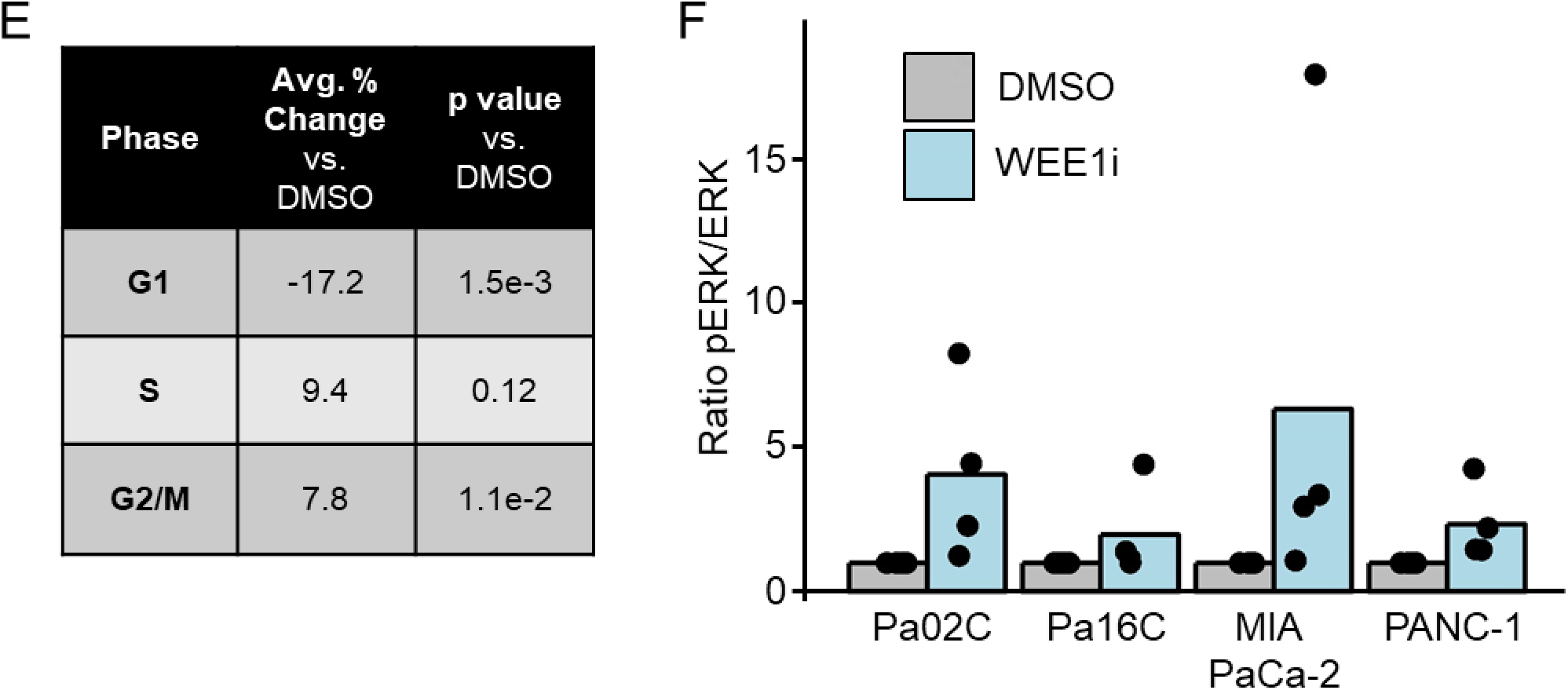
Adavosertib treatment of PDAC induces growth arrest and apoptosis. *A,* immunoblot analysis of PDAC cells following treatment with WEE1i (adavosertib) for 48 h (n = 4). *B,* GI_50_ values were calculated in six PDAC cell lines for the WEE1 inhibitor adavosertib after five days of treatment. Data are representative of three independent experiments. *C,* immunoblot analysis of Pa16C cells following treatment with WEE1i (adavosertib) at various time points (n = 4). *D,* flow cytometry cell cycle analysis was performed following 48 h DMSO or WEE1i (Pa02C 100 nM adavosertib; Pa16C, PANC-1, MIA PaCa-2 200 nM adavosertib) treatment of four PDAC cell lines. Quantification was performed in FCS Express (n = 4). *E,* Tabular summary of results from panel *A*. Student’s *t*-test was used to identify significant differences between adavosertib treatment and DMSO treated samples. *F,* quantification of relative pERK over ERK ratios using densitometry with 100 nM (Pa02C) or 200 nM WEE1i (Pa16C, MIA PaCa-2 and PANC-1) corresponding to (Fig. 4B and Fig. S4*A*) (n = 4).

**Figure S5.**
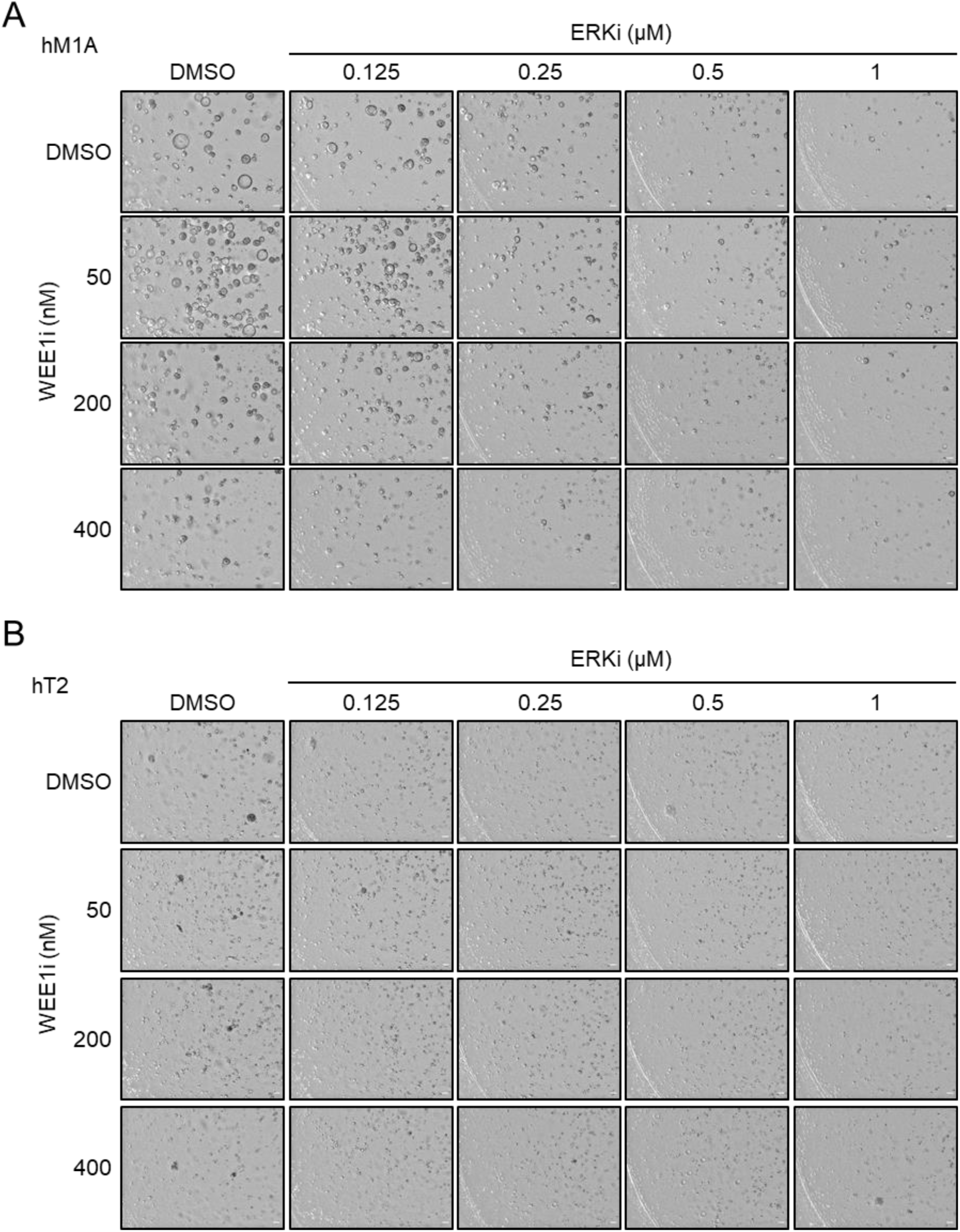
Combined inhibition of WEE1 and ERK inhibits PDAC organoid growth. *A,* hM1A PDAC organoids were seeded for three days in Matrigel and organoid maintenance factors, followed by treatment for 10 days with ERKi or WEE1i alone or in combination across a range of doses. Representative images are shown; scale bar is equivalent to 100 µm (n = 4). *B,* hT2 PDAC organoids were seeded and treated according to the same parameters as hM1A organoids in *A*.

**Table S1.** **KRAS suppression-induced downregulated kinases and their functions.** Kinase functions are adapted from UniProtKB (https://www.uniprot.org/help/uniprotkb) and GeneCards (https://www.genecards.org/)

| Gene | Protein | Function |
| --- | --- | --- |
| *NEK2* | NEK2 | NIMA related kinase 2 Ser/Thr protein kinase involved in mitotic regulation by controlling centrosome separation and bipolar spindle formation, and chromatin condensation in meiotic cells |
| *WEE1* | WEE1 | Ser/Thr protein kinase acts as a negative regulator of entry into mitosis (G2 to M transition) by protecting the nucleus from cytoplasmically activated cyclin B1-complexed CDK1 before the onset of mitosis by mediating phosphorylation of CDK1 |
| *TTK* | TTK | Tyr/Ser/Thr protein kinase essential for chromosome alignment at the centromere during mitosis and is required for centrosome duplication |
| *PLK1* | PLK1 | Polo-like kinase 1 Ser/Thr protein kinase performs several important functions throughout M phase of the cell cycle, including the regulation of centrosome maturation and spindle assembly, the removal of cohesins from chromosome arms, the inactivation of anaphase-promoting complex/cyclosome (APC/C) inhibitors, and the regulation of mitotic exit and cytokinesis |
| *MELK* | MELK | Maternal embryonic leucine zipper Ser/Thr protein kinase involved in cell cycle regulation, mitotic spindle pole assembly, apoptosis and splicing regulation |
| *AURKA* | Aurora A | Ser/Thr protein kinase plays a critical role in microtubule formation and stabilization during chromosome segregation in mitosis |
| *CHEK1* | CHK1 | Ser/Thr protein kinase is required for checkpoint-mediated cell cycle arrest in response to DNA damage. Integrates signals from ATM/ATR protein kinases to facilitate cell cycle inhibition |
| *PKMYT1* | PKMYT1 | Protein kinase, membrane-associated tyrosine/threonine 1 Ser/Thr protein kinase is membrane-associated and negatively regulates the G2/M transition of the cell cycle by phosphorylating CDK1 |
| *EPHA2* | EPHA2 | Ephrin type-A receptor 2 Tyr kinase binds membrane-bound Ephrin-A family ligands, regulating cell adhesion and differentiation; overexpressed in cancer, with tumor promoting role |
| *TNIK* | TNIK | TRAF2 and NKC interacting Ser/Thr protein kinase activates the Wnt-β-catenin signaling pathway, with roles in gene transcription, cytoskeletal rearrangements and cell spreading |
| *CDK1* | CDK1 | Cyclin dependent kinase 1 Ser/Thr protein kinase plays a key role in the control of the eukaryotic cell cycle by modulating the centrosome cycle as well as mitotic onset; promotes G2/M transition and regulates G1 progress and G1/S transition via association with multiple interphase cyclins; required in higher cells for entry into S-phase and mitosis |
| *CDK7* | CDK7 | Cyclin dependent kinase 7 Ser/Thr protein kinase involved in cell cycle control and in RNA polymerase II-mediated RNA transcription; required for activation and complex formation of CDK1/cyclin-B during G2/M transition |
| *CSNK2A1* | CKIIα | Casein kinase 2 alpha 1 Ser/Thr protein kinase involved in various cell cycle control and apoptosis. During mitosis, functions as a component of the p53-dependent spindle assembly checkpoint (SAC) that maintains cyclin-B-CDK1 activity and G2 arrest in response to spindle damage |

**Table S2.**
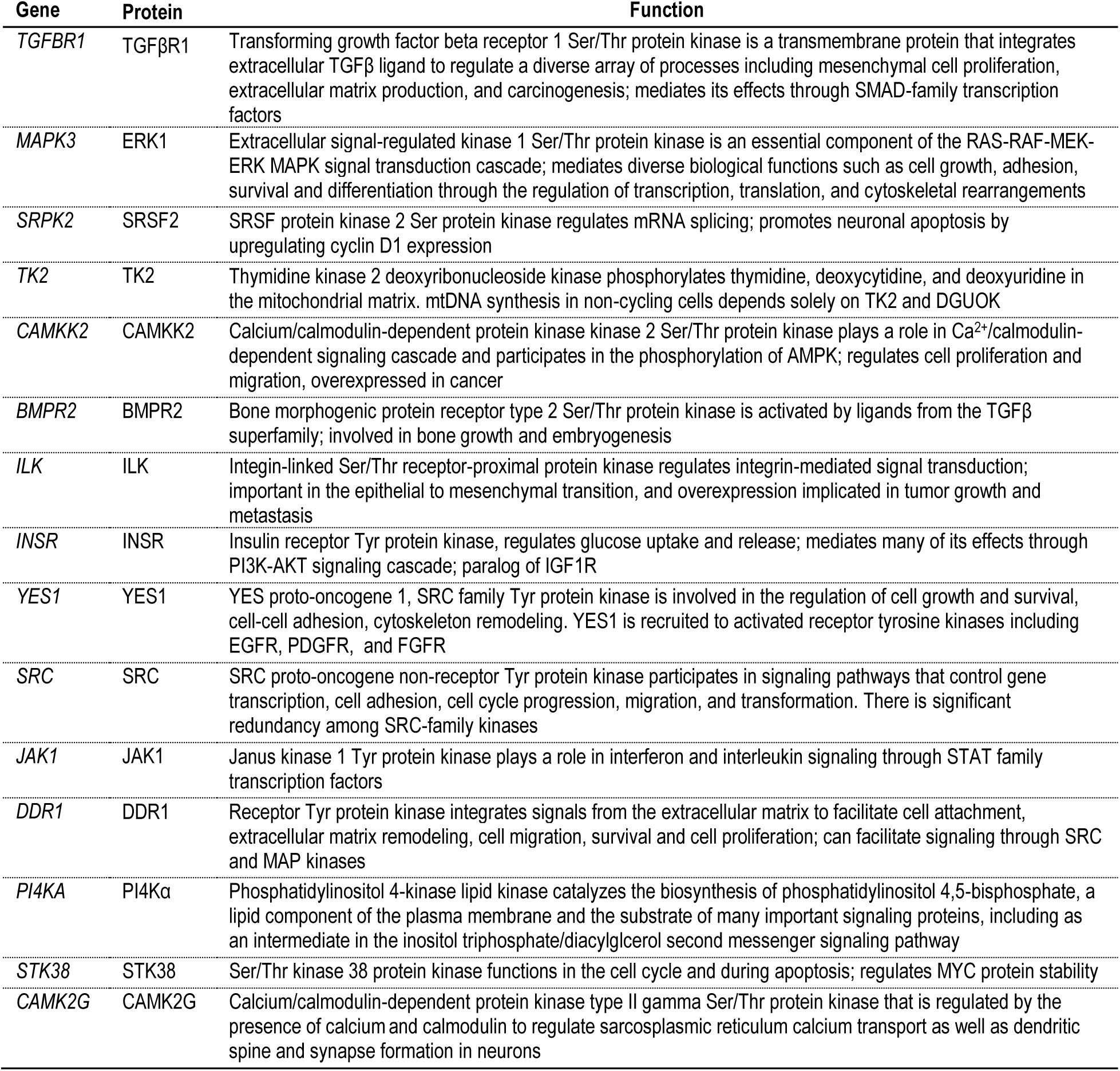
**KRAS suppression-induced upregulated kinases and their functions.** Kinase functions are adapted from UniProtKB (https://www.uniprot.org/help/uniprotkb) and GeneCards (https://www.genecards.org/)

**Table S3.**
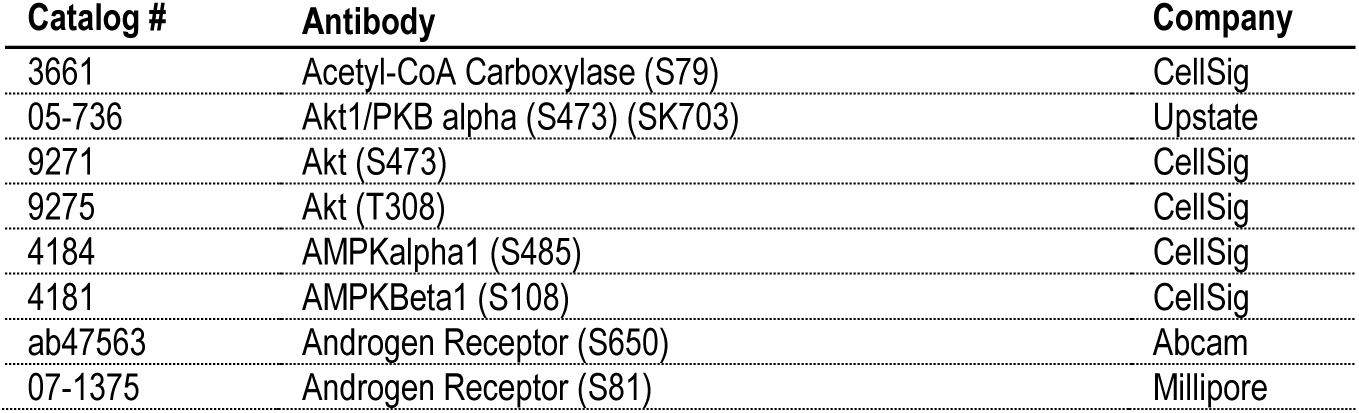
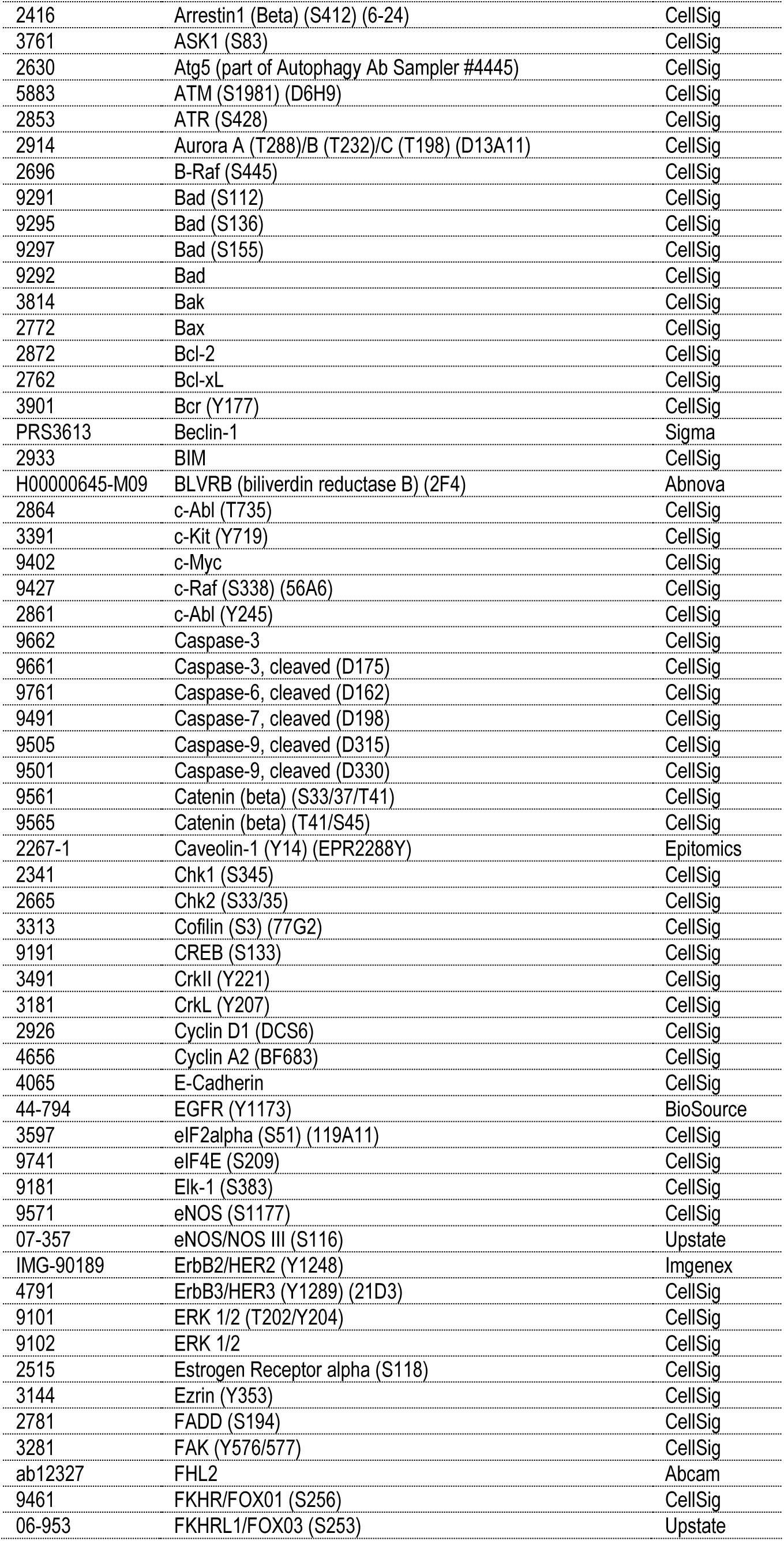
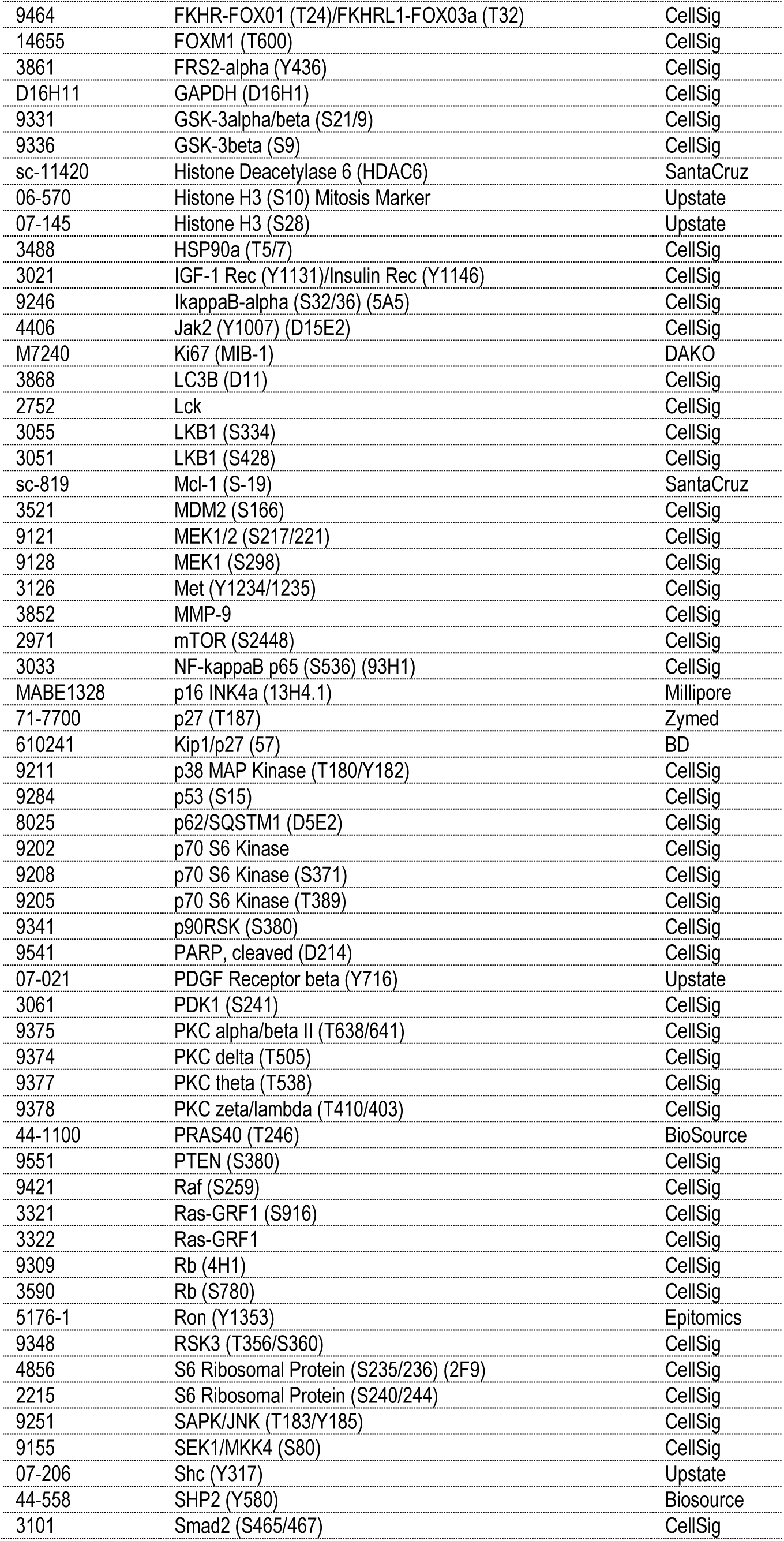
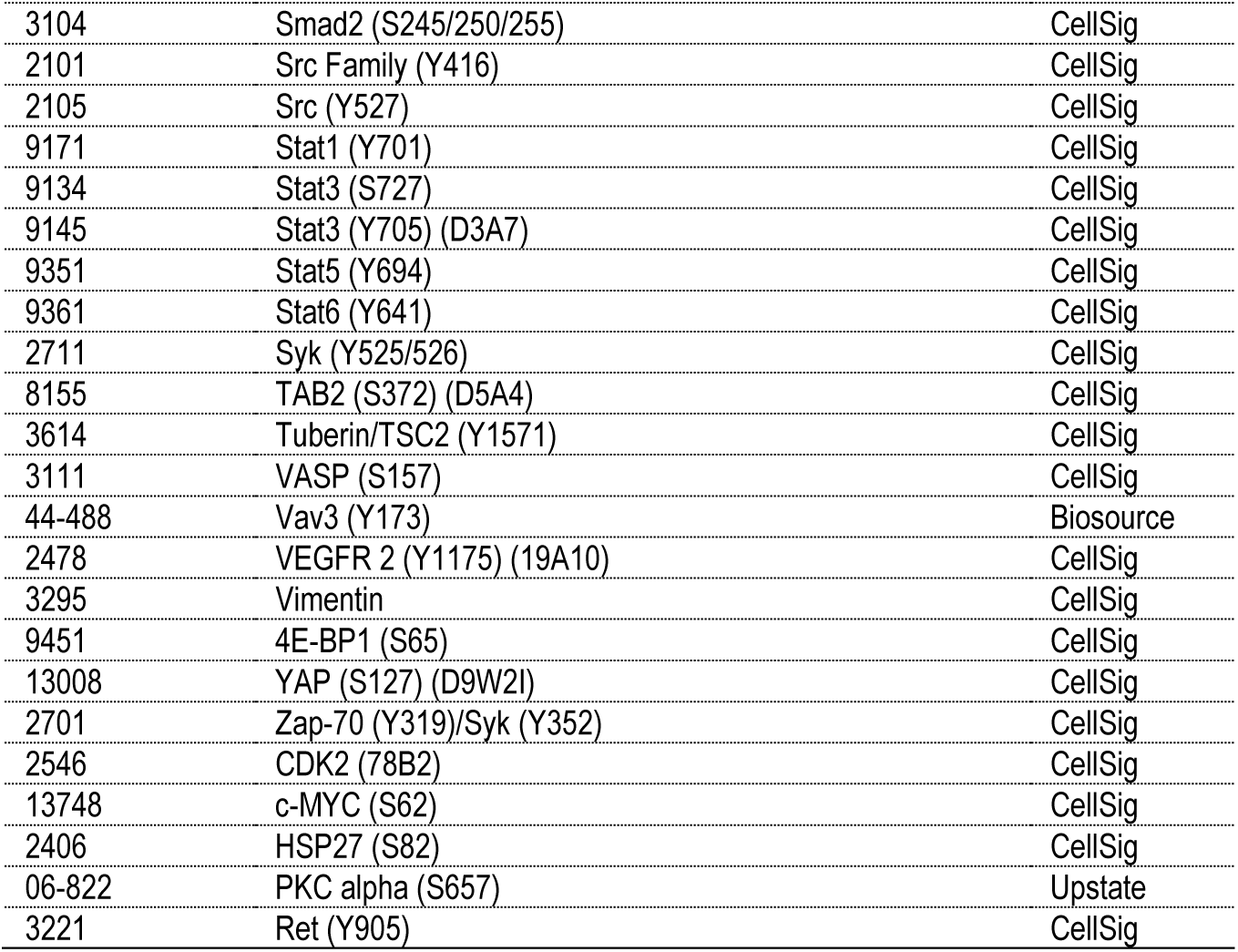
RPPA Antibodies.

